# Neuronal circuitry for stimulus selection in the visual system

**DOI:** 10.1101/598383

**Authors:** António M. Fernandes, Johannes Larsch, Joseph C. Donovan, Thomas O. Helmbrecht, Duncan Mearns, Yvonne Kölsch, Marco Dal Maschio, Herwig Baier

## Abstract

Visual objects naturally compete for the brain’s attention, and selecting just one of them for a behavioural response is often crucial for the animal’s survival^1^. The neural correlate of such stimulus prioritisation might take the form of a saliency map by which responses to one target are enhanced relative to distractors in other parts of the visual field^2^. Single-cell responses consistent with this type of computation have been observed in the tectum of primates, birds, turtles and lamprey^2–7^. However, the exact circuit implementation has remained unclear. Here we investigated the underlying neuronal mechanism presenting larval zebrafish with two simultaneous looming stimuli, each of which was able to trigger directed escapes on their own. Behaviour tracking revealed that the fish respond to these competing stimuli predominantly with a winner-take-all strategy. Using brain-wide functional recordings, we discovered neurons in the tectum whose responses to the target stimulus were non-linearly modulated by the saliency of the distractor. When the two stimuli were presented monocularly in different positions of the visual field, stimulus selection was already apparent in the activity of retinal ganglion cell axons, a likely consequence of antagonistic mechanisms operating outside the classical receptive field^8,9^. When the two stimuli were presented binocularly, i.e., on opposite sides of the fish, our analysis indicates that a loop involving excitatory and inhibitory neurons in the nucleus isthmi (NI) and the tectum weighed stimulus saliencies across hemispheres. Consistent with focal enhancement and global suppression, glutamatergic NI cells branch locally in the tectum, whereas GABAergic NI cells project broadly across both tectal hemispheres. Moreover, holographic optogenetic stimulation confirmed that glutamatergic NI neurons can modulate visual responses in the tectum. Together, our study shows, for the first time, context-dependent contributions of retinotectal and isthmotectal circuits to the computation of the visual saliency map, a prerequisite for stimulus-driven, bottom-up attention.

---

Dark, looming stimuli are strongly aversive stimuli for zebrafish larvae^10,11^ and other animals^12,13^, probably mimicking an approaching predator or an object on a collision course. In our setup, single looming disks presented from below and on one side of a free-swimming animal were highly effective in driving an escape response to the contralateral side (Fig. 1a-c, Extended Data Fig. 1i, j). Depending on the location and the strength of the stimulus, fish larvae adjust direction and magnitude of their response. We identified loom expansion rate and contrast as key factors that modulate escape probability (Extended Data Fig. 1a-e, see also^14^). We asked how zebrafish respond to two looming stimuli presented simultaneously (Fig. 1d). We reasoned that fish may either “select” one of the two stimuli for response and suppress a response to the other stimulus (winner-take-all hypothesis); alternatively, fish might integrate both stimuli, triggering an escape response in a direction along the mean vector of responses to either stimulus presented alone (averaging hypothesis)^15–18^.

**Figure 1.**
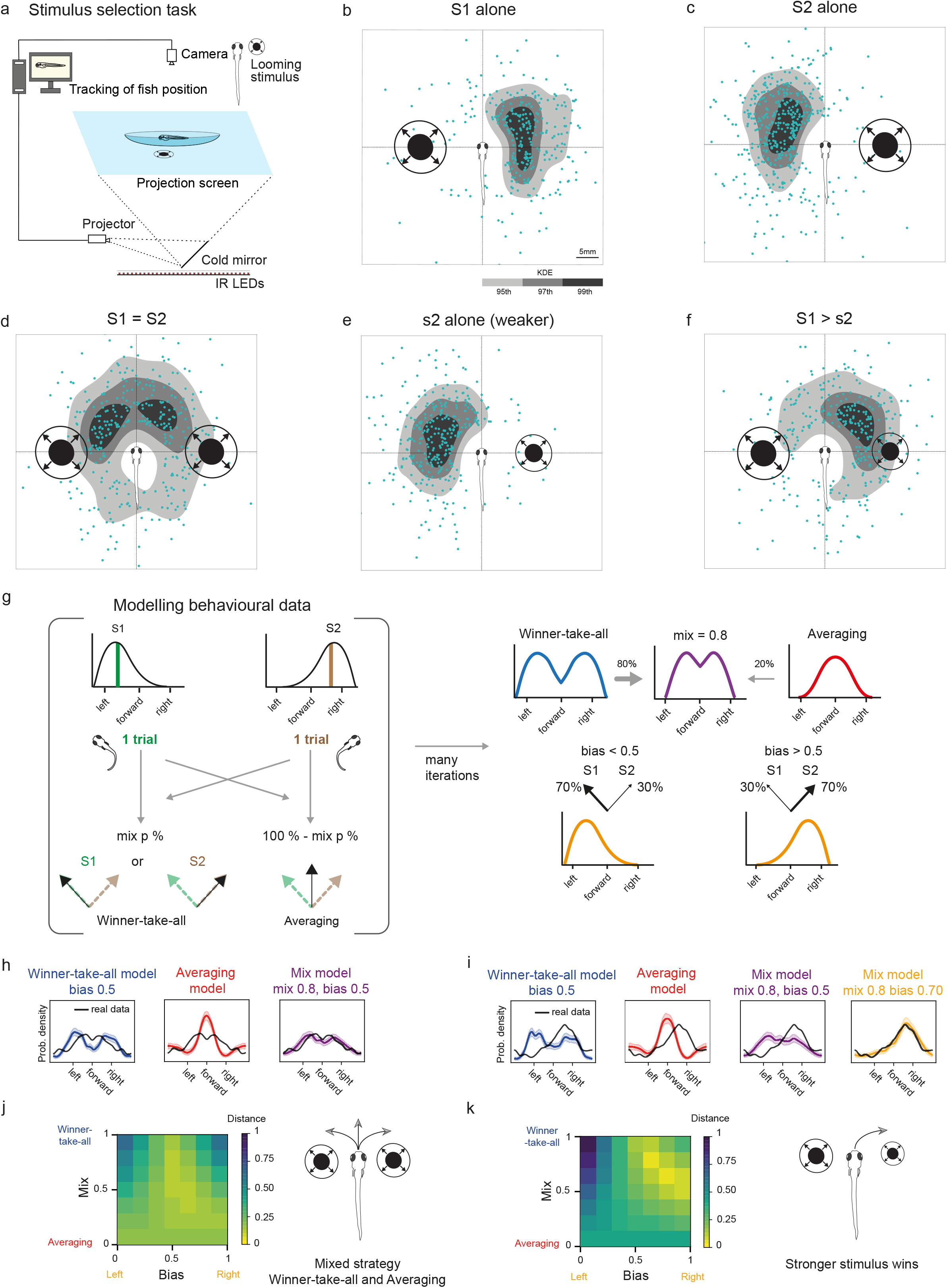
Zebrafish respond to competing stimuli according to their relative saliencies. **a.** Schematic drawing of the behavioural task used for measuring stimulus selection. The animal is tracked while updating in real time the positions of the looming disks projected from below. **b.** Response to a single looming stimulus (S1 alone, 90°/s) presented on the left side of the fish. Blue dots are the XY positions of the fish after escape at the end of the expansion period of the stimulus (500ms). In grayscale are the kernel density estimation (KDE) isocontours of the same data. **c.** Similar to (b), but with stimulus presented on the right side of the fish. **d.** Competition (S1 + S2) of equal stimuli (90°/s). **e.** Weaker stimulus (s2, 60°/s) on the right side of the fish. **f.** Competition (S1 +s2) of unequal stimuli (90°/s vs. 60°/s). **g.** Modelling of the behavioural data, which simulates the distribution of responses to competing stimuli by combing the single trial responses to individual stimuli. One stimulus response from an S1 trial and one stimulus response from an S2 trial are combined using repeated random sampling. The winner-take-all model chooses randomly between the S1 response and the S2 response. The averaging model combines the pair of responses by taking the vector average of the response angle. The mixture model implements a random assortment between the winner-take-all model (with probability p) and the average model (with probability 1-p). **h.** Modelling of behaviour outcome for equal stimuli competition. Shaded areas are 97.5% confidence intervals (CI). **i.** Modelling of behaviour outcome for unequal stimuli competition. Shaded areas are 97.5% CI. **j.** Quality of the behaviour reconstruction. Heatmap showing the normalised energy distance related to the panel (g) depending on the model parameters (Bias and Mix). Bias: represents the probability of response left vs right; Mix: represents the mixing factor between “winner-take-all” and “averaging” models. **k.** Similar to j, but for unequal stimuli. N=117 fish.

When we presented two stimuli of equal strength, appearing on either side of the fish, we observed a bimodal distribution of escape trajectories. Larvae consistently escaped in a sideways direction away from one, apparently randomly chosen disk (Fig. 1d, Extended Data Fig. 1g). Modulating the expansion rate of one stimulus (e.g. slower expansion rate) biased the combined responses away from the stronger stimulus (Fig. 1e-f, Extended Data Fig. 1f-o). While this result argues in favour of a winner-take-all mechanism, a smaller, but significant fraction of responses pointed toward an intermediate direction, consistent with an averaging strategy. To estimate the relative contribution of each strategy, we fit a biased-mixture model implementing predictions from both hypotheses (Fig 1g). For equal stimuli (bias = 0.5), we found that a mix of winner-take-all (80% of responses) and averaging (20% of responses) best explained the data (Fig 1h, j and Extended Data Fig. 1k, m). For unequal stimuli, we additionally fit the bias term (Fig. 1i). This revealed that fish selected the stronger stimulus 70% of the times, (bias term = 0.7) (Fig 1i-k, Extended Data Fig. 1l, n)).

We next asked if the winner-take-all behavioural strategy extended to a situation where two looming stimuli were displayed to the same eye in non-overlapping parts of the visual field (Extended Data Fig. 2). A single looming disk, positioned in the posterior visual field, triggered a forward escape, (47° +/- 3.9 SEM), whereas an anteriorly located disk triggered a sideways escape (82.5° +/- 7.6 SEM) (Extended Data Fig. 2l). Both stimuli together triggered a distribution of escape angles that included the responses to single stimuli. The limited dynamic range of escape angles for the two stimuli precluded fitting our biased-mixture model. However, as with binocular stimulation, the faster of two monocular stimuli dominated escape direction such that its mean angle was indistinguishable from that triggered by the stimulus alone (Extended Data Fig. 2l). Thus, stimulus selection is also detectable with monocular stimuli.

Next we investigated the potential neural correlates of stimulus selection using brain-wide calcium imaging (Fig. 2a, b). We first determined which regions responded reliably to looming stimuli (Extended Data Fig. 3a). As shown previously^10,19^, looming stimuli activated retinal ganglion cell (RGC) axons, the tectum, the pretectum and a thalamic area near retinal arborisation field AF4. We also found a responsive area at the midbrain-hindbrain boundary that we identified as the putative zebrafish homolog of the nucleus isthmi (NI)^20^, a region that has previously been implicated in the generation of a visual ‘saliency map’^3,6,21–23^.

**Figure 2.**
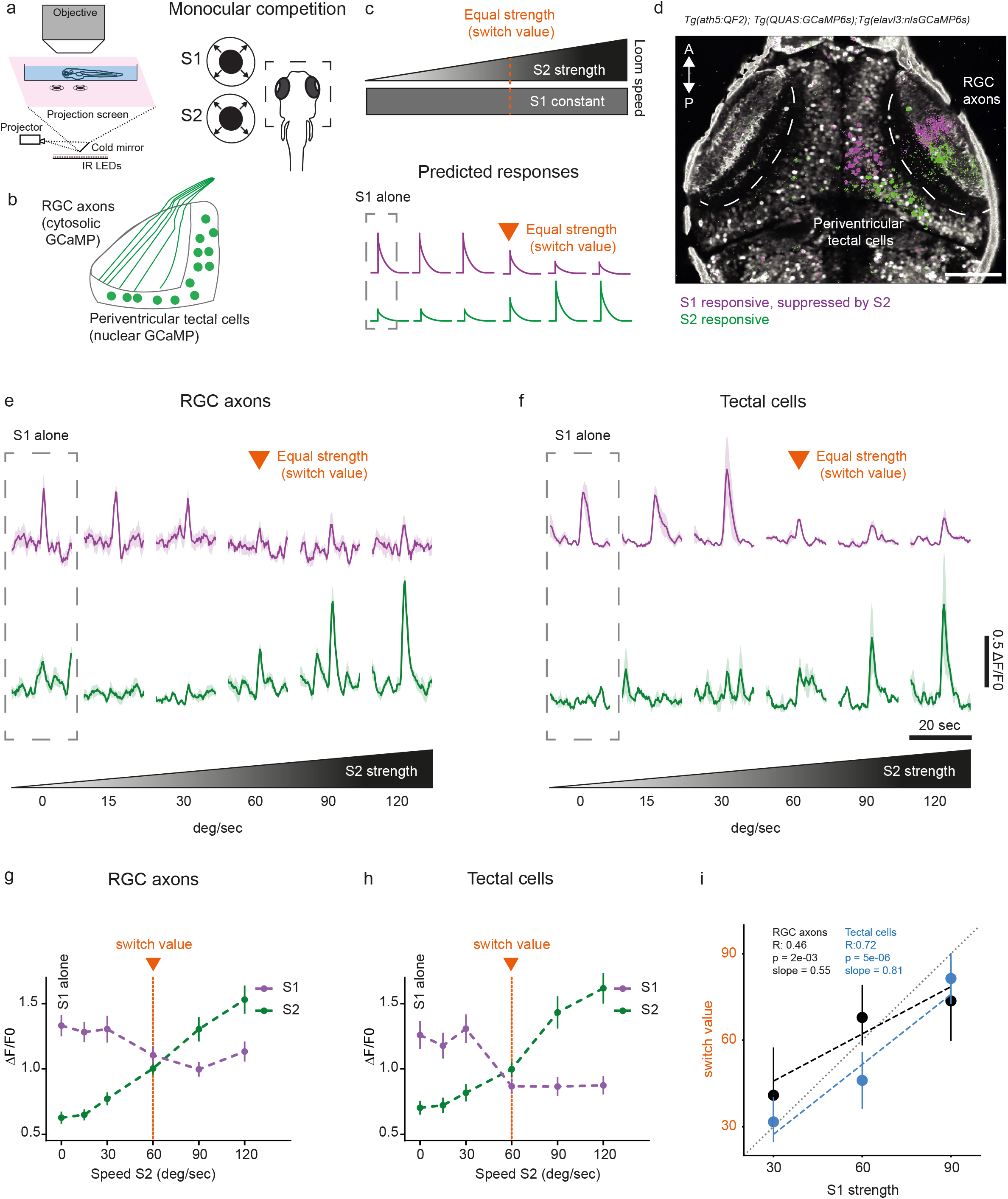
Retinal ganglion cell and tectal activity is suppressed during monocular competition. **a.** Left panel, schematic drawing of calcium imaging setup. Right panel, monocular competition task. S1: stimulus 1, located in the anterior part of the visual field. S2: stimulus 2, located in the posterior part of the visual field. **b.** Schematic of the expression pattern of triple-transgenic fish used for this experiment *(Tg(ath5:QF2; QUAS:GCaMP6s; elavl3:nlsGCaMP6s))*. Simultaneous recording of the activity of RGC axons and tectal cells was carried out by combining an RGC-specific enhancer (*ath5:QF2*), driving expression of cytoplasmic GCaMP6s (*QUAS:GCaMP6s*) (see Extended Data Fig. 9), and a pan-neuronal enhancer, driving nuclear-localised GCaMP6s (*elavl3:nlsGCaMP6s*). **c.** Schematic of competition protocol. Expansion rate of one looming stimulus is kept constant (S1), while systematically varying the velocity of the competitor stimulus (S2). In orange, condition where both stimuli have equal strength (“switch value”). Below is shown the predicted responses types accounting for stimulus selection **d.** Pixel-wise regression analysis of the temporal series during a single imaging trial. Corresponding t-statistic for each pixel is calculated. Map shows associated S1-responsive pixels, suppressed by a stronger S2 stimulus (in magenta), and pixels that enhance their responses as a function of S2 strength (in green). Scale bar represents 50 μm. **e.** Characteristic activity profiles for RGCs. Top traces, average of 10 RGC axon ROIs that were suppressed by a stronger S2 stimulus (in magenta). Lower traces, average of 10 RGC axon ROIs that were enhanced by S2 (in green). **f.** Characteristic activity profiles for tectal periventricular neurons. Top traces, average of 10 tectal ROIs that were suppressed by a stronger S2 stimulus (in magenta). Lower traces, average of 10 tectal ROIs that were enhanced by S2 (in green). S2 strength is indicated below. Orange arrow shows condition where both stimuli have equal strength (“switch value”). **g.** Summary plot across all conditions for RGC axon pixels. Switch-like responses, showing RGC pixels suppressed by S2, are shown in magenta. RGC pixels that were enhanced by S2 are shown in green. **h.** Summary plot across all conditions for tectal pixels. Switch-like responses, showing tectal pixels suppressed by S2, are shown in magenta. Tectal pixels that were enhanced by S2 are shown in green. **i.** Switch value increases with S1 strength for both RGC axons and tectal cells. R-value is the correlation coefficient and the p-value relates to testing whether the slope is zero. N=5 fish. Errors are SEM.

Competing ensembles of tectal neurons have been observed in the zebrafish tectum^24^. We hypothesised that we should find at least two response types to the competing looming response: (i) neurons whose activities scale with the strength of one stimulus and (ii) neurons whose activities are suppressed by the competing stimulus. We designed a protocol to find these two response types. We kept the expansion rate of one looming stimulus constant (S1), while systematically varying the velocity of the competitor stimulus (S2) (Fig. 2c). Presenting two competing stimuli to the same eye resulted in the suppression of activity in a subset of tectal cells (Fig. 2d, f, h, in magenta). The response of these cells to S1 was substantially reduced, when S2 was stronger or identical to S1 (Fig. 2f) but was high when S2 was weaker than S1. On the other hand, we found responses that scaled with increasing S2 speed (Fig. 2d, f, h, in green). These findings are consistent with stimulus competition by reciprocal inhibition^25,26^. Similar response profiles to looming stimuli have previously been called “switch-like” in the barn owl^22^. Remarkably, we observed switch-like responses already at the level of the RGC axons (Fig. 2d, e, g, Extended Data Fig. 3b-c). This suggests that monocular stimulus competition affects the activity of RGCs and may be inherited by tectal cells. In agreement with previous reports^22,25^, the switch transition for the population response is flexible and shifts systematically with the strength of the S1 stimulus (Fig 2i, Extended Data Fig. 3c, e). Tectal cells are more switch-like compared to RGCs, suggesting that saliency computation is amplified in the tectum (correlation coefficients, RGCs: R = 0.47, Tectum: R = 0.72, Fig. 2i). To ask if the stimulus competition extends to stimuli with different valence, we designed synthetic, prey-like stimuli, which evoke hunting behavior^27^. Indeed, RGC axons and tectal responses showed suppression and enhancement driven by a competing prey stimulus on the same hemisphere (Extended Data Fig. 4a-c). Such a mechanism might facilitate efficient target selection during hunting against a background of distractors.

The suppression observed in RGC terminals is likely the result of intraretinal processing of competing stimuli by means of lateral inhibition^28,29^. To rule out that RGC axon terminals receive feedback modulation within the tectum^30^, we ablated the tectal cells and then imaged the RGC terminals in response to competing looming stimuli. Switch-like responses of RGCs remained intact upon removal of tectal influences (Extended Data Fig. 4d-h). Whereas responses in RGCs appeared unaffected, tectal ablation led to severe impairments in responses to prey and looming stimuli as reported previously^10,27,31^ (Extended Data Fig. 4i-4l). These results indicate that the formation of saliency maps previously attributed to computations in higher-order visual areas already begins in the inner retina.

Two stimuli presented to opposite sides of the fish (Fig. 3a), produced switch-like suppression and enhancement of distinct populations of neurons, similar to same-eye stimulation (Fig. 3c). We, therefore, predicted the existence of a circuit that compares signals between the two eyes. Such dynamics were observed both in the tectum and the NI across hemispheres (Fig. 3b-g). Using transmitter-specific lines^32^, we determined that, as in other vertebrate species, cells in the NI either express glutamatergic (*vglut2a*) or GABAergic (*gad1b*) markers in a mutually exclusive fashion and express known marker genes for the isthmic region (e.g. Reelin)^33^ (Fig. 3h, Extended Data Fig. 5). Some of the glutamatergic cells are also cholinergic. The glutamatergic/cholinergic and GABAergic populations form two spatially segregated nuclei across the hindbrain-midbrain boundary (Fig. 3h). Functional recordings revealed that both NI populations display switch-like activity profiles; i. e., the response to the target stimulus was suppressed by a competitor stimulus. However, only the glutamatergic neuronal activity scales with the strength of the competitor (Extended Data Fig. 6).

**Figure 3.**
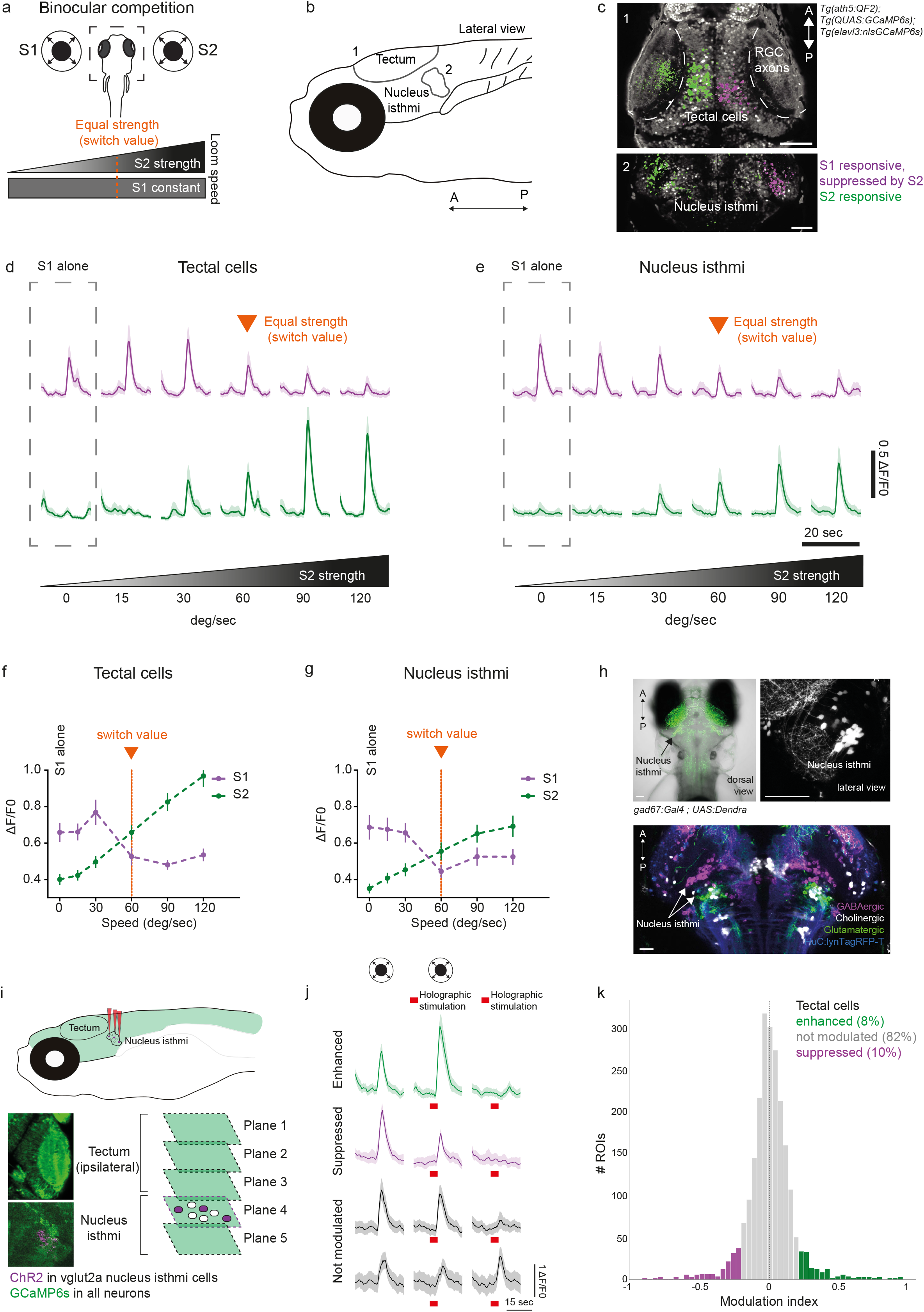
Winner-take-all dynamics in tectal and isthmic neurons in response to competing binocular stimuli. **a.** Binocular competition task. S1: stimulus 1, located on the left side of the fish. S2: stimulus 2, located on the right side of the fish. In orange, condition where both stimuli have equal strength (“switch value”). **b.** Anatomical location of the tectum and the nucleus isthmi (NI) in zebrafish larvae. **c.** Pixel-wise regression analysis of the temporal series during a single imaging trial. The t-statistic for each pixel is calculated. Map 1 shows associated S1-responsive tectal pixels, suppressed by a stronger S2 stimulus (in magenta). Tectal pixels that enhance their response as a function of S2 intensity are shown in green. Map 2 shows the same response profiles as in Map 1 but for the nucleus isthmic. Scale bars represent 50 μm. **d.** Characteristic activity profiles for tectal periventricular neurons. Top traces, average of 10 tectal ROIs that were suppressed by a stronger S2 stimulus (in magenta). Lower traces, average of 10 tectal ROIs that were enhanced by S2 (in green). **e.** Characteristic activity profiles for NI neurons. Top traces, average of 10 NI ROIs that were suppressed by a stronger S2 stimulus (in magenta). Lower traces, average of 10 NI ROIs that were enhanced by S2 (in green). S2 strength is indicated below. Orange arrow shows the condition where both stimuli have equal strength (“switch value”). **f.** Summary plot across all conditions for tectal pixels. Switch-like responses, showing tectal pixels suppressed by S2, are shown in magenta. Tectal pixels that were enhanced by S2 are shown in green. N=5 fish. **g.** Summary plot across all conditions for NI pixels. Switch-like responses showing NI pixels that were suppressed by S2 are shown in magenta. NI pixels that were enhanced by S2 are shown in green. N=4 fish. h. Top left panel shows a dorsal image of a double-transgenic *Tg(gad1b: Gal4VP16)mpn155; Tg(UAS:Dendra-kras)s1998t* fish, labelling GABAergic neurons in green. Arrow indicates location of GABAergic NI neurons. Top right panel shows lateral view of *Tg(gad1b: Gal4VP16)mpn155; Tg(UAS:nfsb-mCherry)c264* fish, labeling GABAergic neurons in white. Lower panel shows the alignment of transgenic lines used to label selectively the NI populations. *Tg(gad1b: Gal4VP16)mpn155*, labeling GABAergic NI neurons (magenta), *Tg(lhx9: Gal4VP16)mpn203*, labeling glutamatergic NI neurons (green) and *Tg(chata:Gal4VP16)mpn202*, labeling cholinergic NI neurons (white). *Tg(elavl3:lyn-tagRFP)mpn404* is used as a reference channel (blue). Scale bars represent 50 μm. **i.** Schematic of the experiment during two-photon computer-generated holography (2P-CGH) activation of specific excitatory isthmic neurons expressing channelrhodopsin (ChR2), combined with volumetric imaging of ipsilateral tectal responses. **j.** Photostimulation of excitatory isthmic neurons modulates tectal responses during visual stimulation (responses to looming). Some of the tectal cell responses are unaffected by optogenetic stimulation (in grey), while others are either suppressed (in magenta) or enhanced (in green). **k.** Histogram showing quantification of tectal response modulation. Modulation index is defined as ((visual alone) - (visual combined with optogenetic stimulation)) / ((visual alone) + (visual combined with optogenetic stimulation)). N=4 fish.

To test whether there is a functional connection between the NI and tectum, we imaged tectal and isthmic activity in response to a looming stimulus, while optogenetically activating a subpopulation of excitatory NI neurons (Fig. 3i). We observed a strong modulation of the stimulus-evoked activity in the contralateral NI and tectum; responses were either enhanced or suppressed upon co-activation of glutamatergic NI neurons (Fig. 3j, k, Extended Data Fig. 7). This suggests that photostimulation likely activates different functional classes of neurons in the NI. As in adult frogs^34^, activation of NI neurons alone (without a visual stimulus present) did not reliably lead to activation of tectal cells (Extended Data Fig.7a-d).

Intertectal and tectobulbar projection neurons were recently described in larval zebrafish^35^. Anatomical co-registration revealed that tectal axon terminals overlap with dendritic arborisations of NI neurons on both sides of the brain (Fig. 4a, b, Extended Data Fig. 8b-d). To search for isthmo-tectal connections, we stochastically labelled single NI cells and traced their projections (Fig. 4c-f, Extended Data Fig. 8a-o). In agreement with our optogenetic results, we found two classes of excitatory NI neurons innervating both hemispheres along different pathways (Fig. 4c, d, Extended Data Fig. 8k-l). Glutamatergic axons arborise focally in either the ipsilateral or the contralateral tectum. In contrast, GABAergic neurons arborise broadly in either one or both tecta (Fig. 4e, f, Extended Data Fig. 8i-j, 8n). This anatomical architecture is in agreement with our binocular competition results in the NI. In addition, it supports previous findings showing interocular interactions between left and right NI^36^. Thresholding of the difference in activity of left and right tectum has been proposed as a possible mechanism to compare stimulus saliency in mammals^37^.

**Figure 4.**
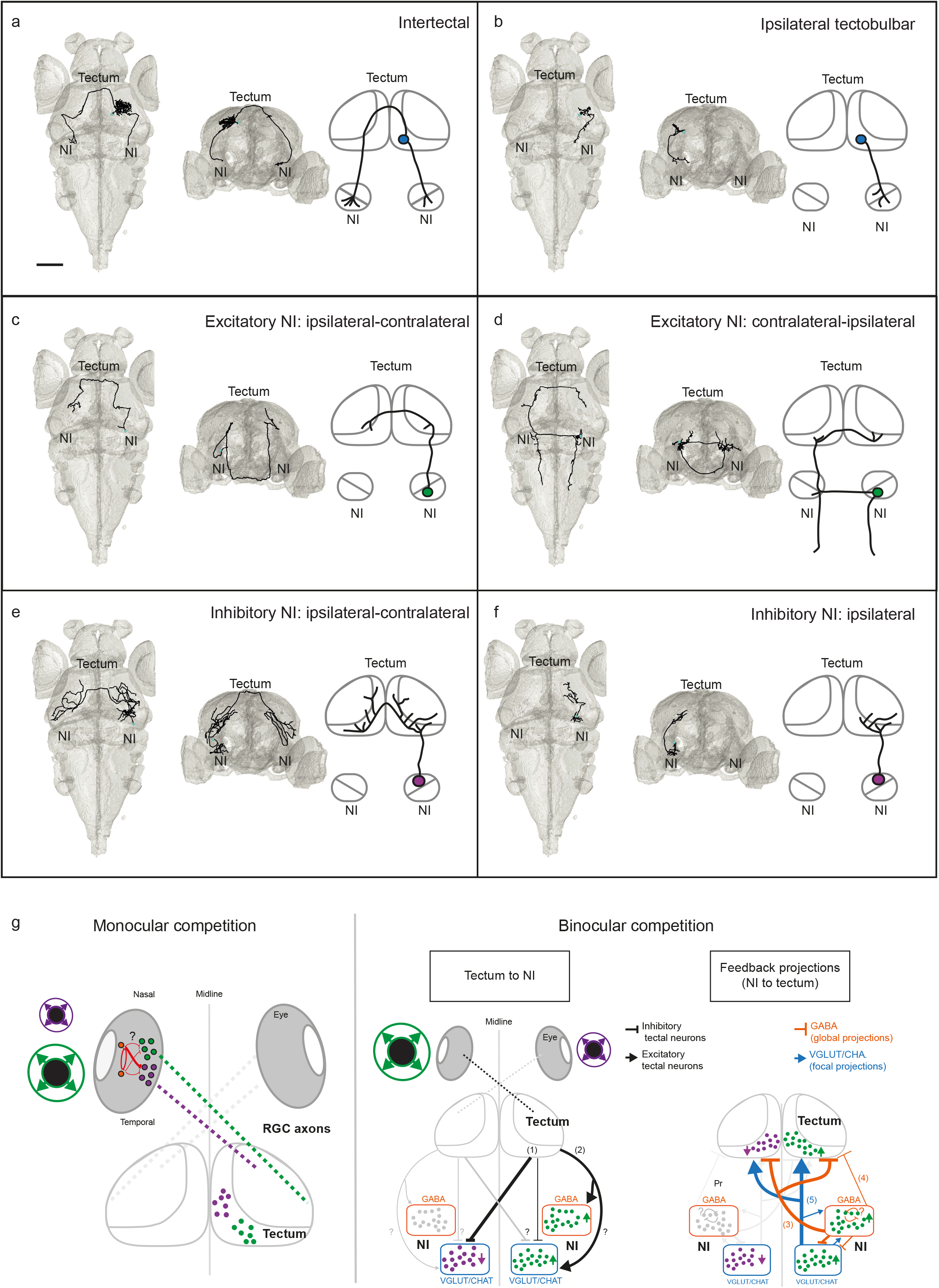
The isthmotectal loop is a possible substrate for binocular competition. **a-f.** Single cell neuronal reconstructions (black traces). For each morphological type, two views are shown (anterior-posterior and medial-lateral axis), plus a schematic of isthmotectal circuitry (right). Scale bar represents 50 μm. **g.** Summary of findings. Left: Hypothetical model for monocular competition. Amacrine cells (orange) inhibit each other and suppress post-synaptic RGCs (magenta). RGCs that respond to the most salient, “winning” stimulus, are highly active (green). The result of this competition is conveyed to the tectum through RGC axons and further augmented by a tectum-intrinsic circuit. Right: Anatomical connectivity of the isthmotectal loop and hypothetical circuit model for binocular competition. Tectal cells are depicted in black. Putative inhibitory intertectal cells form axon collaterals to (1) differentially inhibit excitatory NI cells on the ipsilateral and contralateral side. Putative excitatory tectal projection neurons (2) activate both inhibitory and excitatory NI neurons on the ipsilateral side. Intertectal cells project mainly to the dendrites of excitatory NI cells (Extended Data Fig. 8d). Ipsilateral tectal projection neurons terminate in the excitatory and inhibitory neuropil of the ipsilateral NI (Extended Data Fig. 8b-c). Selection of most salient stimulus is done across the hemispheres. “Winning” stimulus activates both contralateral inhibitory and contralateral excitatory NI neurons (green). “Losing” the competition leads to suppression of the excitatory NI population (magenta). Feedback projections from the NI to the tectum are shown in orange (inhibitory) and blue (excitatory). Reciprocal projections between the excitatory and inhibitory NI cells are shown inside the NI box. GABAergic NI neurons project via a superficial commissure and arborise broadly in the contralateral and ipsilateral tecta (3) or only the ipsilateral tectum (4), where they may implement reciprocal inhibition across hemispheres. Excitatory NI neurons cross the midline via the postoptic commissure, located deep in the diencephalon. One class of cells form collaterals in both the ipsilateral and the contralateral tectum (5) (see Extended Data Fig. 8k), where they enhance the winning activity (green cells in the tectum). Suppressed tectal cells are shown in magenta. The other class of excitatory NI cells projects first to the contralateral glutamatergic NI, with arborisations close to the pretectum, thalamus and a neuropil region close to the contralateral semicircular torus and tectum, and then returns to the ipsilateral side s (Fig. 4d, Extended Data Fig. 8f, 8h and 8l). We posit that this delayed excitation may balance the system, once the behavioural response is finished. The third class of excitatory cells projects only to the ipsilateral thalamus (Extended Data Fig. 8e, m). Question marks highlight circuit components whose neurotransmitter identity or connections are unknown. NI: Nucleus isthmi. See also Extended Data Fig. 8.

In conclusion, the topographical arrangement and transmitter identities of recurrent connections in the isthmotectal loop support a saliency map mechanism, in which representation of one stimulus is focally enhanced, while responses to stimuli elsewhere are suppressed (Fig. 4g). Such a network could produce the observed winner-take-all outcome of behavioural choice during stimulus competition^25,38,26^. Together, a feed-forward retinotectal and a modulatory isthmotectal recurrent circuit implement context-dependent target selection (Fig. 4g) and might form the basis of a bottom-up attentional mechanism.

## Acknowledgements

The authors thank Anna Krammer, Yunmin Wu, David Northmore, Aristides Arrenberg and Ruben Portugues for technical advice and critical feedback. Enrico Kühn, Styliani Koutsouli, Krasimir Slanchev and Irene Arnold-Ammer provided technical support. Vilim Štih and Andreas Kist shared scripts used in the analysis of imaging data. K. Kawakami provided the *SAGFF(LF)81C* zebrafish line. W. Driever provided the riboprobes for *gad67* and *trh*. We thank Eva Laurell, Michael Kunst and Nouwar Mokayes for help with experiments and analysis of some of the cells used for the anatomical reconstruction data. We thank all members of the Baier lab for critical discussions and helpful comments. Funding was provided by the Max Planck Society and the DFG (SFB 870 “Assembly and Function of Neural Circuits” and Priority Programme SPP 1926 “Next Generation Optogenetics”).

## Author Contributions

A.M.F. and H.B. conceived the project. A.M.F. designed and performed experiments and analyzed the data. J.L. build the behavioural setup and performed some of the behavioural data analysis. T.O.H. wrote some of the analysis code for imaging experiments and anatomical reconstruction data. Y.K. generated new lines and expression data for Q-system genetic tools used in the paper. D.M. performed the experiments and analysis regarding prey capture data. J.C.D. performed the modelling analysis of behavioural data and wrote parts of the code used to analyze imaging data. M.D.M. performed some of the experiments and analysis regarding imaging and optogenetic results. A.M.F. and H.B. wrote the manuscript with the assistance of all authors. All authors discussed the data and the manuscript.

