## Extended data for "Neuronal circuitry for stimulus selection in the visual system"

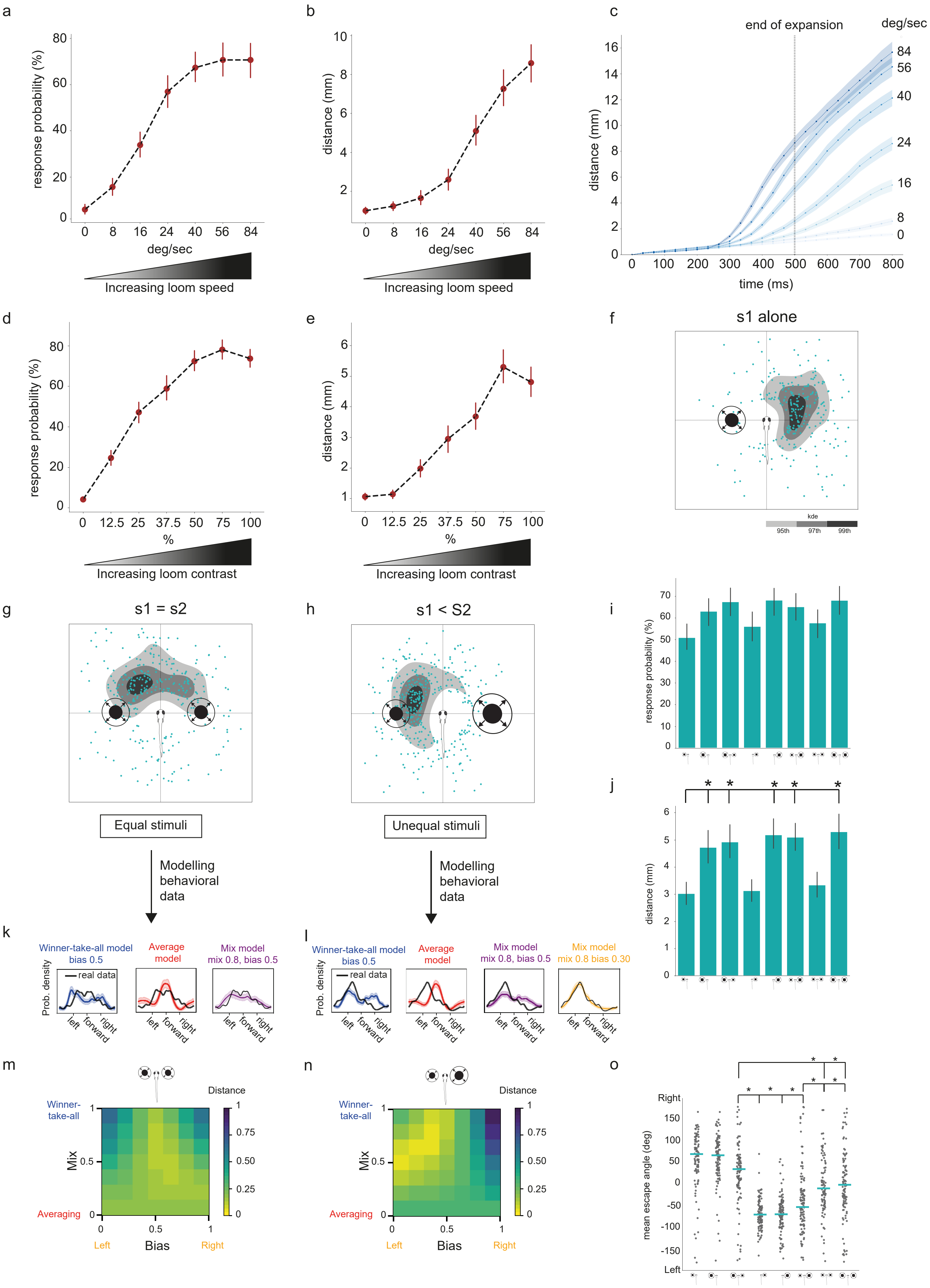

a
response probability (%)
deg/sec
Increasing loom speed
b
distance (mm)
deg/sec
Increasing loom speed
c
end of expansion
deg/sec
distance (mm)
time (ms)
d
response probability (%)
%
Increasing loom contrast
e
distance (mm)
%
Increasing loom contrast
f
s1 alone
kde
95th
97th
99th
g
s1 = s2
Equal stimuli
Modelling behavioral data
h
s1 < S2
Unequal stimuli
Modelling behavioral data
i
response probability (%)
j
distance (mm)
k
Winner-take-all model bias 0.5
Average model
Mix model mix 0.8, bias 0.5
real data
Prob. density
left
forward
right
l
Winner-take-all model bias 0.5
Average model
Mix model mix 0.8, bias 0.5
Mix model mix 0.8 bias 0.30
real data
Prob. density
m
Distance
Winner-take-all
Mix
Averaging
Bias
Left
Right
n
Distance
Winner-take-all
Mix
Averaging
Bias
Left
Right
o
Right
Left
mean escape angle (deg)

#### <u>Extended Data</u>

**Extended Figure 1.** Zebrafish modulate their escape behaviour in response to differences in velocity and contrast of looming stimuli.

**a, b.** Probability of escape (a) and distance travelled (b) increase with looming velocity (expansion rate). **c.** Time course plot of distances travelled under different velocity conditions. Vertical line marks the end of the expansion (500 ms) for stimuli. **d, e.** Probability of escape (d) and distance travelled (e) increase with stimulus contrast. **f.** Response to a single looming stimulus (s1 alone, 60°/s) presented on the left side of the fish. Blue dots are the XY positions of the fish following their escape at the end of the expansion period of the stimulus (500ms, 60°/s). In grayscale are the kernel density estimation (KDE) isocontours of the same data. **g.** Competition (s1 +s2) of equal stimuli (60°/s). **h.** Competition (s1 +S2) of unequal stimuli (60°/s vs. 90°/s). **i., j.** Escape probability (i) and distance travelled (j) for all binocular competition conditions. **k.** Modelling of behaviour outcome for equal stimuli competition (60°/s). Shaded areas are 97.5% confidence intervals (CI). **l.** Modelling of behaviour outcome for unequal stimuli competition (60°/s vs. 90°/s). Shaded areas are 97.5% CI. **m.** Heatmap showing the normalised energy distance from panel (k) depending on the model parameters (Bias and Mix). Bias: represents the probability of response left vs right; Mix: represents the mixing factor between “winner-take-all” and “averaging” models. **n.** Heatmap showing the normalised energy distance from panel (l) depending on the model parameters (Bias and Mix). **o.** Summary plot showing mean escape angle for all conditions during binocular competition. * marks significant comparisons (p-value < 0.05). N=117 fish.

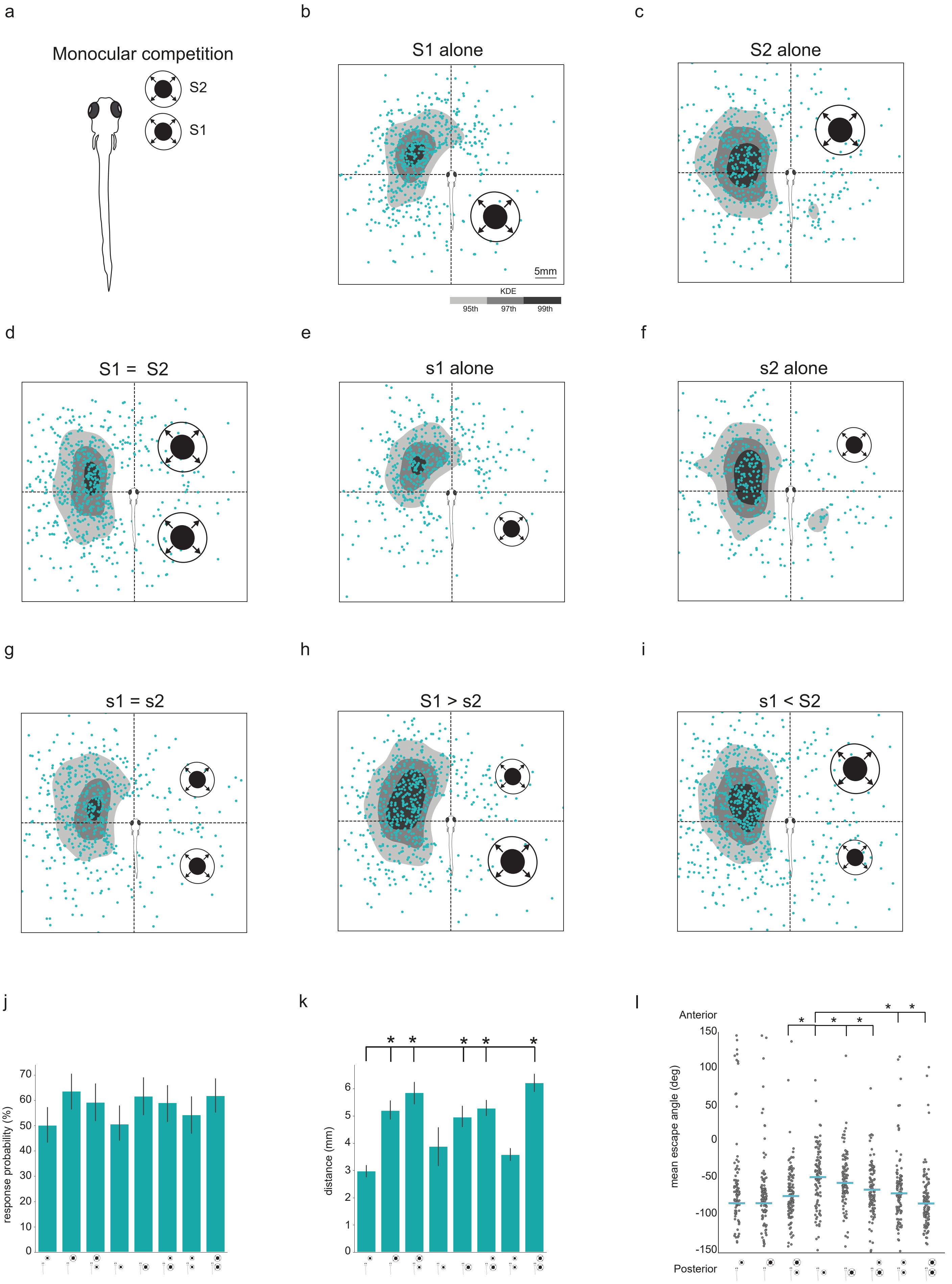
a
Monocular competition
S2
S1
b
S1 alone
5mm
KDE
95th
97th
99th
c
S2 alone
d
S1 = S2
e
s1 alone
f
s2 alone
g
s1 = s2
h
S1 > s2
i
s1 < S2
j
response probability (%)
k
distance (mm)
l
Anterior
Posterior
mean escape angle (deg)

**Extended Figure 2.** Monocular stimulus competition.

**a.** Schematic of monocular competition paradigm. Two stimuli are presented in the visual field of the same eye, posteriorly and anteriorly at 135° or 45°, relative to the fish centre of mass, respectively, with 0 being the fish's initial heading direction. **b.** Response to looming stimulus (S1 alone, 90°/s) presented posteriorly. Blue dots are the XY positions of the fish at the end of the expansion period of the stimulus (500ms) after escape responses elicited by the stimulus. In grayscale, the kernel density estimation of the blue dots, respectively corresponding to $95^{th}$, $97^{th}$ and $99^{th}$ percentile. **c.** Response to looming stimulus (S2 alone, 90°/s) presented anteriorly. **d.** Competition (S1 +S2) of equal strength stimuli (90°/s). **e.** Response to looming stimulus (s1 alone, 60°/s) presented on the posterior part of the visual field. **f.** Response to looming stimulus (s2 alone, 60°/s) presented on the anterior part of the visual field. Competition (s1 +s2) of equal strength stimuli (60°/s). **h.** Competition (s1 +S2) of unequal stimuli (60°/s vs. 90°/s). **i.** Competition (S1 +s2) of unequal stimuli (90°/s vs. 60°/s). **j.** Response probability plot for all monocular competition conditions. **k.** Distance travelled by the fish for all monocular competition conditions. **l.** Summary plot showing mean escape angles for all conditions during monocular competition. * marks significant comparisons (*p*-value < 0.05). N=126 fish.

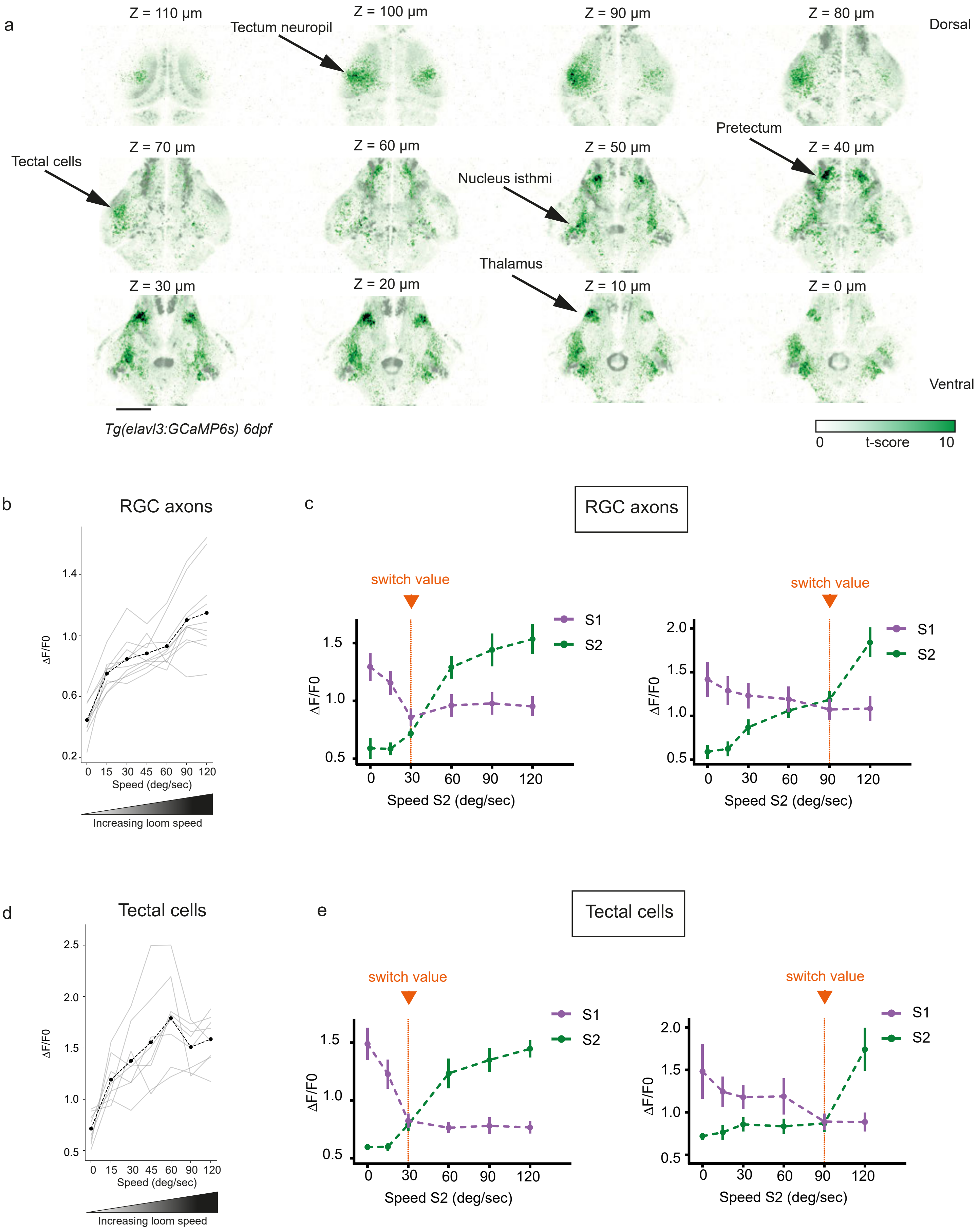

a
Z = 110 µm
Z = 100 µm
Z = 90 µm
Z = 80 µm
Dorsal
Tectum neuropil
Pretectum
Tectal cells
Z = 70 µm
Z = 60 µm
Z = 50 µm
Z = 40 µm
Nucleus isthmi
Thalamus
Z = 30 µm
Z = 20 µm
Z = 10 µm
Z = 0 µm
Ventral
Tg(elavl3:GCaMP6s) 6dpf
0 t-score 10
b
RGC axons
ΔF/F0
Speed (deg/sec)
Increasing loom speed
c
RGC axons
switch value
S1
S2
ΔF/F0
Speed S2 (deg/sec)
d
Tectal cells
ΔF/F0
Speed (deg/sec)
Increasing loom speed
e
Tectal cells
switch value
S1
S2
ΔF/F0
Speed S2 (deg/sec)

**Extended Figure 3.** Whole-brain imaging identifies regions active during stimulus competition.

**a.** Whole-brain imaging of regions that respond reliably to looming stimuli (dark green). Scale bar is 100 μm. **b.** RGC axon activity increases with faster looming velocity. Strong stimuli drive RGC axons to saturation. N=3 fish. **c.** Tectal cell activity increases with faster looming velocity. Strong stimuli drive tectal cells to saturation. N=2 fish. **d.** Summary plot across all conditions for RGC axon pixels. Suppressed RGC pixels are shown in magenta. Enhanced RGC pixels are shown in green (left pane: S1 expansion rate is 30°/s; right panel: S1 expansion rate is 90°/s). N=5 fish. **e.** Summary plot across all conditions for tectum pixels. The response function shows switch-like behaviour. * marks significant comparisons (*p*-value $< 0.05$, Tukey's HSD pairwise test).

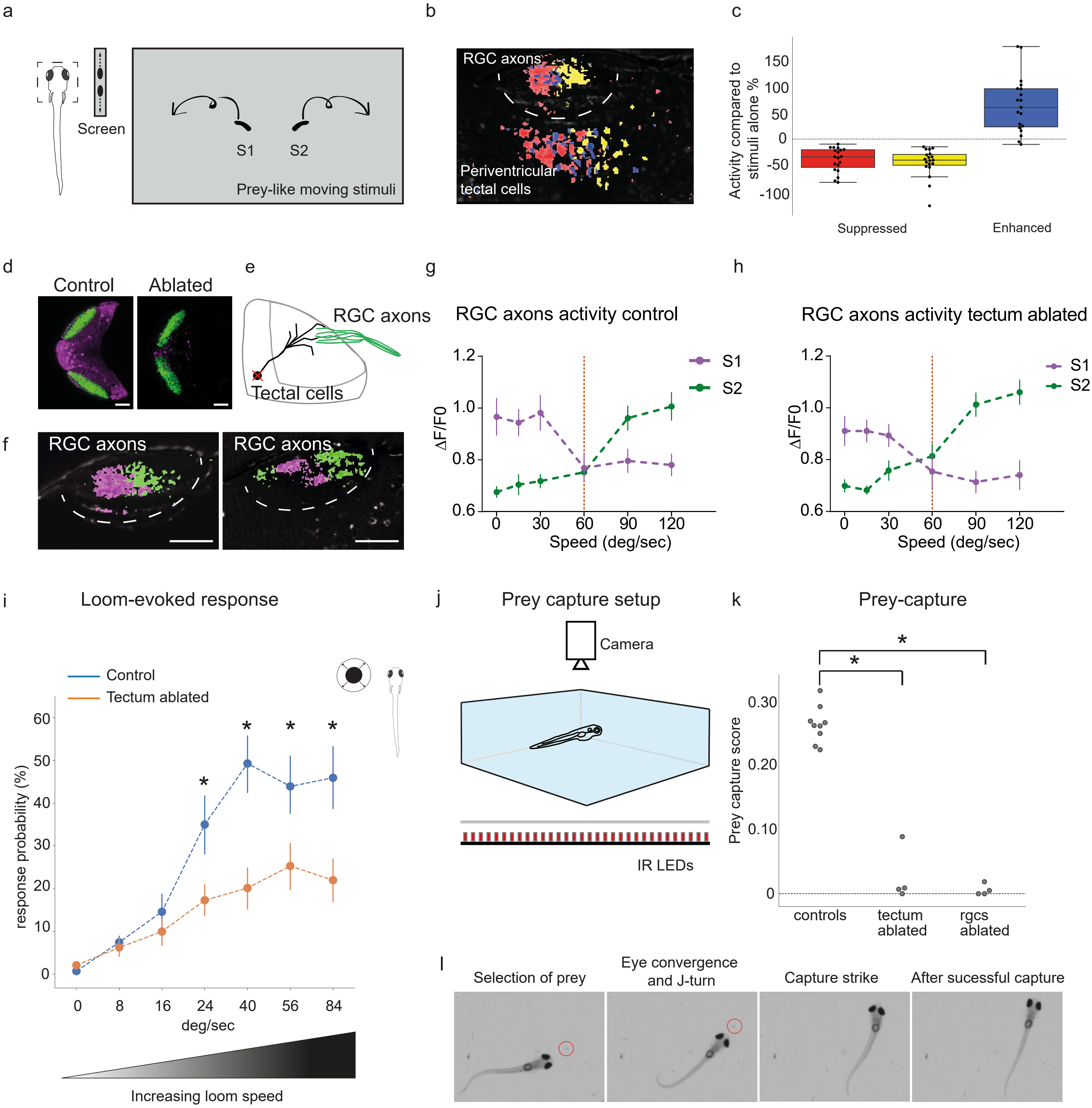
a
Screen
S1
S2
Prey-like moving stimuli
b
RGC axons
Periventricular tectal cells
c
Activity compared to stimuli alone %
150
100
50
0
-50
-100
Suppressed
Enhanced
d
Control
Ablated
e
RGC axons
Tectal cells
f
RGC axons
RGC axons
g
RGC axons activity control
ΔF/F0
1.2
1.0
0.8
0.6
0
30
60
90
120
Speed (deg/sec)
S1
S2
h
RGC axons activity tectum ablated
i
Loom-evoked response
Control
Tectum ablated
response probability (%)
0
8
16
24
40
56
84
deg/sec
Increasing loom speed
j
Prey capture setup
Camera
IR LEDs
k
Prey-capture
Prey capture score
0.30
0.20
0.10
0
controls
tectum ablated
rgcs ablated
l
Selection of prey
Eye convergence and J-turn
Capture strike
After sucessful capture

**Extended Figure 4.** Suppression of activity by competitor stimulus is detectable in RGC axons and tectal cells.

**a.** Monocular competition paradigm for prey-like moving stimuli. **b.** Presentation of competing prey-like stimuli in the monocular visual field leads to suppression (red pixels responding to S1 and yellow pixels responding to S2) and enhancement of responses (blue pixels respond to both stimuli, with overlapping receptive fields for S1 and S2) in RGC axons and tectal cells. Scale bar is 50 µm. **c.** Quantification of responses to competing prey-like stimuli. Activity was normalised to activity in response to a single stimulus. N=5 fish. **d.** Chemogenetic ablation of tectal cells does not affect suppression observed in RGC axons (*p*=0.1797, two-way Mann Whitney). Genotype used was quadruple-transgenic: *ath5:QF2*; *QUAS:GCaMP6s* (green); *SAGFF(LF)81C*; *UAS:NTR-mCherry* (magenta). Left: Control fish. Right: Ablated fish. Scale bars are 100 µm. **e.** Schematic of the experiment. Ablation of tectal cells removes potential modulatory feedback connections from tectum to RGC axons. **f.** Pixelwise regression analysis of the temporal series during a single imaging trial. Corresponding *t*-statistic for each pixel is calculated. Map shows associated S1-responsive pixels, suppressed by a stronger S2 stimulus (in magenta) and pixels that show enhanced responses as a function of S2 strength (in green). Scale bars are 50 µm. Left panel: Control fish. Right panel: Ablated fish. **g.** Summary plot across all conditions for RGC axon pixels in control fish. N=4 fish. **h.** Summary plot across all conditions for RGC axon pixels in ablated fish. N=5 fish. Switch-like responses showing suppressed RGC pixels are shown in magenta. Enhanced RGC pixels are shown in green (S1 expansion rate is 60°/s, right panel). **i.** Ablation of tectal cells leads to impaired responses to looming stimuli (control in blue, ablated fish in orange). N=45 fish. * marks significant comparisons (*p*-value < 0.05). **j.** Schematic of the setup used to record prey capture events in larval zebrafish. **k.** Ablation of tectal cells leads to impaired responses to prey items (paramecia). Shown is prey capture score for control fish (non-expressors treated with MTZ, n=9 fish), tectum ablated fish (*SAGFF(LF)81C; UAS:NTR-mCherry*, treated with MTZ, n=4 fish) and RGC-ablated fish (*ath5:Gal4; UAS:NTR-mCherry*, treated with MTZ, n=4 fish). Control vs. RGC ablation: *p*<0.05, control vs. tectum ablation: *p*<0.05, RGC ablation vs. tectum ablation: p=0.689, Mann-Whitney-U followed by Bonferroni's correction. * marks significant comparisons (*p*-value<0.05). **l.** Example of hunting sequence leading to the successful capture of paramecia (red circle).

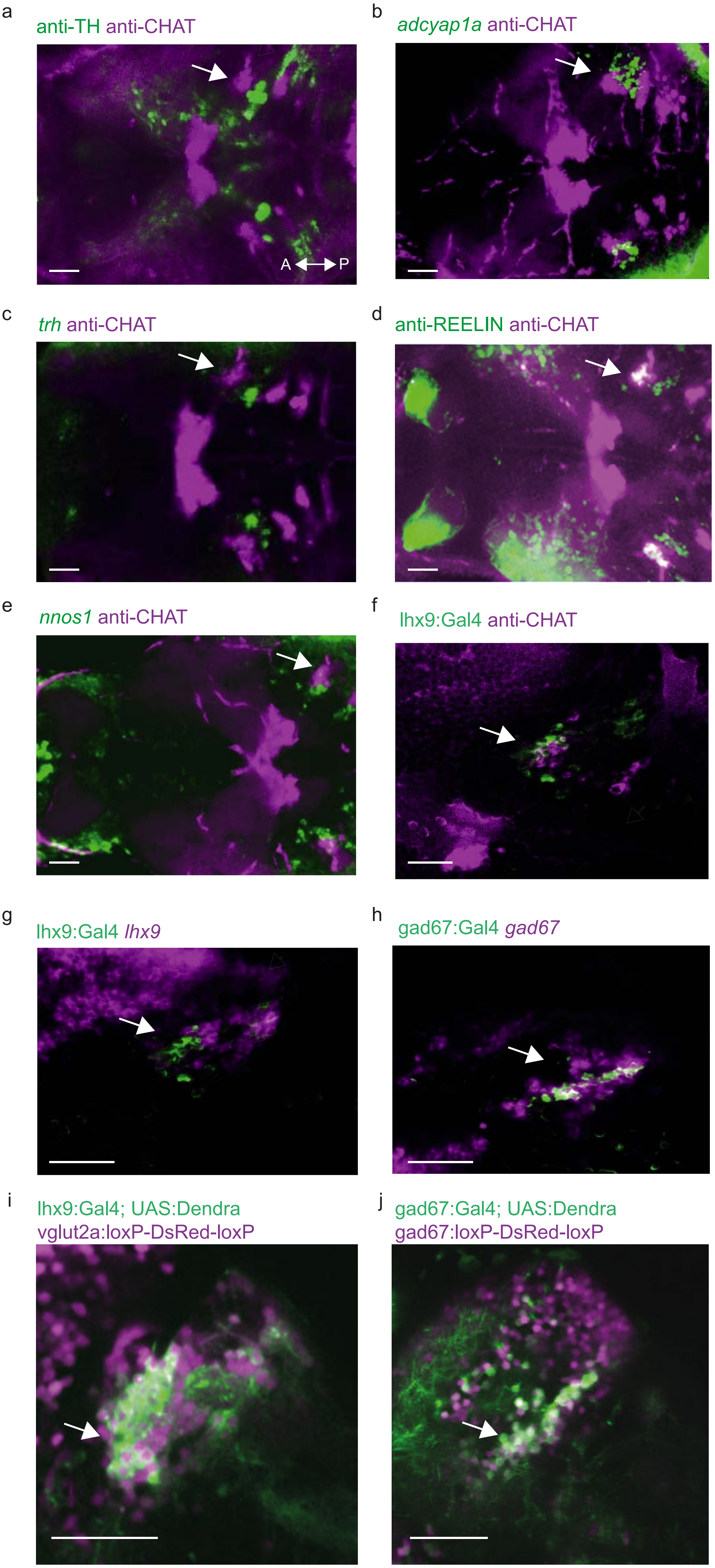

a
anti-TH anti-CHAT
A P
b
adcyap1a anti-CHAT
c
trh anti-CHAT
d
anti-REELIN anti-CHAT
e
nnos1 anti-CHAT
f
lhx9:Gal4 anti-CHAT
g
lhx9:Gal4 lhx9
h
gad67:Gal4 gad67
i
lhx9:Gal4; UAS:Dendra
vglut2a:loxP-DsRed-loxP
j
gad67:Gal4; UAS:Dendra
gad67:loxP-DsRed-loxP

**Extended Figure 5.** Nucleus isthmi is evolutionarily conserved in larval zebrafish.

**a.** Tyrosine hydroxylase (TH) immunohistochemistry labelling (in green) showing Locus Coeruleus (LC) in the vicinity of cholinergic nucleus isthmi cells (arrow points to nucleus isthmi, anti-ChAT in magenta). **b.** *adcyap1a* expression (in green) in the vicinity of cholinergic nucleus isthmi cells (in magenta, white arrow). **c.** *Thyrotropin-releasing hormone (trh)* (in green) in the vicinity of cholinergic nucleus isthmi cells (in magenta, white arrow). **d.** REELIN immunohistochemistry labelling (in green) showing colocalisation with cholinergic nucleus isthmi cells (in magenta, white arrow). **e.** *Nitric oxide synthase (nnos1)* expression (in green) in the vicinity of cholinergic nucleus isthmi cells (in magenta, white arrow). **f.** lhx9:Gal4 line expression (in green) showing colocalisation with cholinergic nucleus isthmi cells (in magenta, white arrow). **g.** lhx9:Gal4 line expression (in green) showing colocalisation with *lhx9* expression in the nucleus isthmi (in magenta, white arrow). **h.** gad67:Gal4 line expression pattern (in green) showing colocalisation with *gad67* mRNA expression in the nucleus isthmi (in magenta, white arrow). **i.** lhx9:Gal4; UAS:Dendra live expression (in green) showing colocalisation with *vglut2a* neurons in vglut2a:loxP-DsRed-loxP line in the nucleus isthmi (in magenta, white arrow). **j.** gad67Gal4; UAS:Dendra live expression (in green) showing colocalisation with *gad67* neurons in gad67:loxP-DsRed-loxP line in the nucleus isthmi (in magenta, white arrow). Scale bars represent 25 μm.

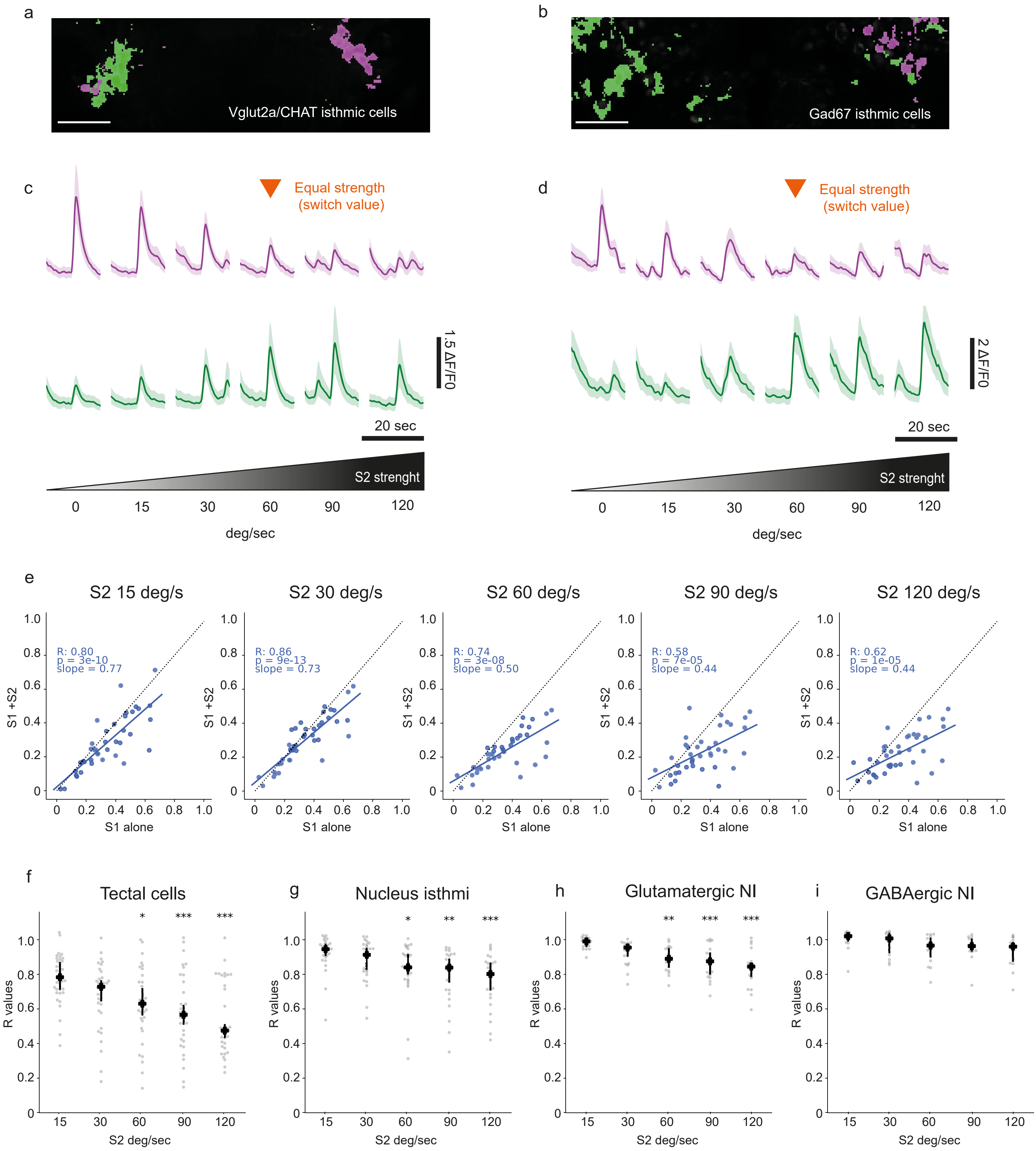

a
Vglut2a/CHAT isthmic cells
b
Gad67 isthmic cells
c
Equal strength
(switch value)
1.5 ΔF/F0
20 sec
S2 strenght
0
15
30
60
90
120
deg/sec
d
Equal strength
(switch value)
2 ΔF/F0
20 sec
S2 strenght
0
15
30
60
90
120
deg/sec
e
S2 15 deg/s
R: 0.80
p = 3e-10
slope = 0.77
S2 30 deg/s
R: 0.86
p = 9e-13
slope = 0.73
S2 60 deg/s
R: 0.74
p = 3e-08
slope = 0.50
S2 90 deg/s
R: 0.58
p = 7e-05
slope = 0.44
S2 120 deg/s
R: 0.62
p = 1e-05
slope = 0.44
S1 +S2
S1 alone
f
Tectal cells
g
Nucleus isthmi
h
Glutamatergic NI
i
GABAergic NI
R values
S2 deg/sec
15
30
60
90
120
*
**
***

**Extended Figure 6.** Glutamatergic but not GABAergic nucleus isthmi cells show significant suppression by competing stimulus across the hemispheres.

**a.** Map shows associated S1 responsive glutamatergic (*lhx9:Gal4; UAS:GCaMP6s* positive) nucleus isthmi pixels, suppressed by a stronger S2 stimulus (in magenta). Pixels that enhance their response as a function of S2 intensity are shown in green. **b.** Map shows associated S1 responsive gabaergic (*gad1b:Gal4; UAS:GCaMP6s* positive) nucleus isthmi pixels, suppressed by a stronger S2 stimulus (in magenta). Pixels that enhance their response as a function of S2 intensity are shown in green. Scale bars represent 25 µm. **c.** Average of 10 selected glutamatergic nucleus isthmi ROIs suppressed by a stronger S2 stimulus (in magenta). Lower traces, average of 10 selected nucleus isthmi ROIs enhanced by S2 strength (in green). Below is shown schematically S2 strength. Orange arrow shows condition where both stimuli have equal strength. **d.** Average of 10 selected gabaergic nucleus isthmi ROIs suppressed by a stronger S2 stimulus (in magenta). Lower traces, average of 10 selected gabaergic nucleus isthmi ROIs enhanced by S2 strength (in green). Below is shown schematically S2 strength. Orange arrow shows condition where both stimuli have equal strength. **e.** Single-trial example of correlation of activity of nucleus isthmi ROIs (elavl3:nlsGCaMP6s positive cells in the NI) between trials with single stimuli and competition trials. X-axis shows stimulus alone (S1, 60°/s) and Y-axis shows the activity of ROIs during different competition conditions (from left to right, S2 competitor stimulus with 15, 30, 60, 90 and 120 deg/s expansion rate). **f.** Summary plot for correlation (R-values for each trial is shown in grey and mean + 95% CI is shown in cyan) across multiple conditions for tectal ROIs (elavl3:nlsGCaMP6s tectal positive cells). N=5 fish. **g.** Summary plot for correlation (R-values for each trial is shown in grey and mean + 95% CI is shown in cyan) across multiple conditions for nucleus isthmi ROIs (elavl3:nlsGCaMP6s NI positive cells). N=4 fish. **h.** Summary plot for correlation (R-values for each trial is shown in grey and mean + 95% CI is shown in cyan) across multiple conditions for glutamatergic nucleus isthmi ROIs (*lhx9:Gal4; UAS:GCaMP6s* positive cells). N=2 fish. **i.** Summary plot for correlation (R-values for each trial is shown in grey and mean + 95% CI is shown in cyan) across multiple conditions for GABAergic nucleus isthmi ROIs (*gad1b:Gal4; UAS:GCaMP6s* positive cells). N=2 fish. Asterisks mark significant comparisons (* p-value < 0.05, (** p-value < 0.01, (*** p-value < 0.001).

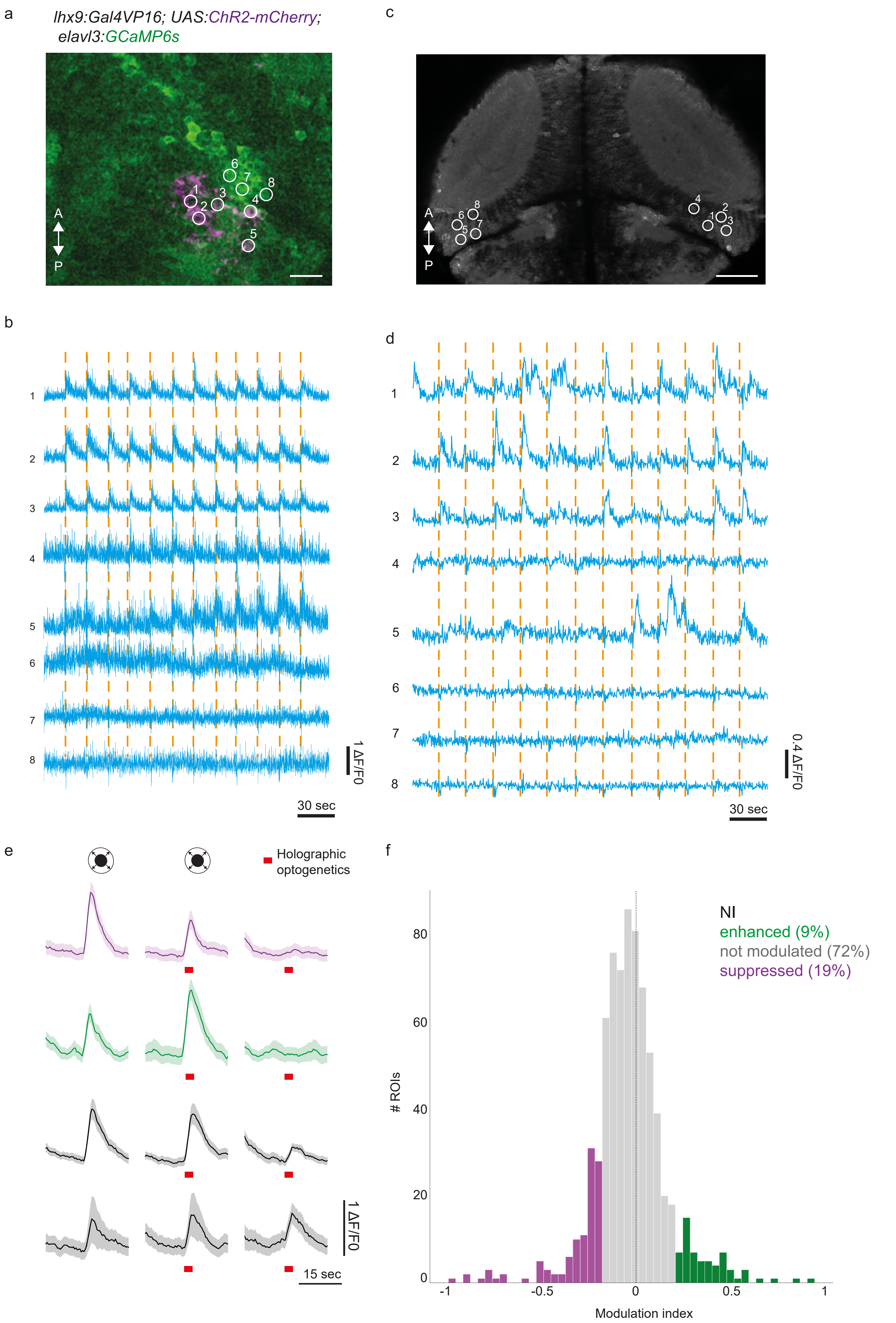
a
lhx9:Gal4VP16; UAS:ChR2-mCherry;
elavl3:GCaMP6s
A
P
c
A
P
b
1 ΔF/F0
30 sec
d
0.4 ΔF/F0
30 sec
e
Holographic optogenetics
1 ΔF/F0
15 sec
f
NI
enhanced (9%)
not modulated (72%)
suppressed (19%)
### ROIs
Modulation index

**Extended Figure 7.** Holographic optogenetics activation of glutamatergic nucleus isthmi population

**a.** Expression of pan-neuronal cytosolic calcium indicator (GCaMP6s, in green) and ChR2-mCherry in glutamatergic nucleus isthmi cells (in magenta). Scale bar represents 15 µm. **b.** Optogenetic activation of glutamatergic nucleus isthmi neurons induced by 1000 ms photostimulation at 920 nm while imaging at 1,020 nm. **c.** Expression of pan-neuronal cytosolic calcium indicator (GCaMP6s, in grey) in both tecta. Selected ROIs are shown. Scale bar represents 50 µm. **d.** Activity of ROIs in (c) driven by optogenetic stimulation of glutamatergic nucleus isthmi neurons. **e.** Photostimulation of excitatory isthmic neurons modulates isthmic region responses during visual stimulation (responses to looming). Some of the isthmic ROIs responses are unaffected by optogenetic stimulation (in grey), while others are either suppressed (in magenta) or enhanced (in green). **f.** Summary histogram showing quantification of modulation of isthmic responses. Modulation index is defined as ((visual alone) - (visual combined with optogenetic stimulation)) / ((visual alone) + (visual combined with optogenetic stimulation)). N=4 fish.

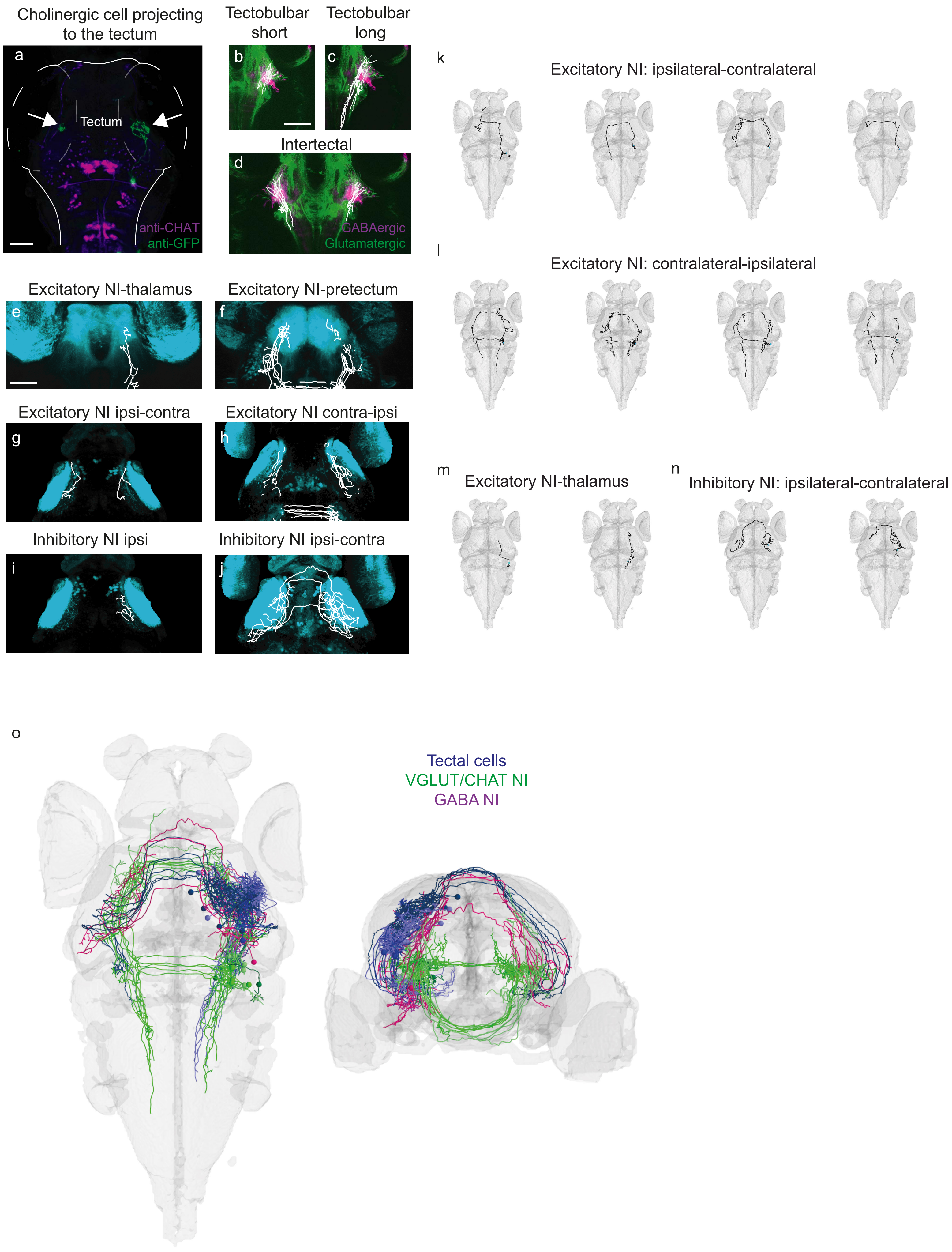

Cholinergic cell projecting to the tectum
a
Tectum
anti-CHAT
anti-GFP
Tectobulbar short
b
Tectobulbar long
c
Intertectal
d
GABAergic
Glutamatergic
Excitatory NI-thalamus
e
Excitatory NI-pretectum
f
Excitatory NI ipsi-contra
g
Excitatory NI contra-ipsi
h
Inhibitory NI ipsi
i
Inhibitory NI ipsi-contra
j
k
Excitatory NI: ipsilateral-contralateral
l
Excitatory NI: contralateral-ipsilateral
m
Excitatory NI-thalamus
n
Inhibitory NI: ipsilateral-contralateral
o
Tectal cells
VGLUT/CHAT NI
GABA NI

**Extended Figure 8.** Nucleus isthmi forms a feedback loop with the tectum

**a.** Example of sparse labelling of a cholinergic nucleus isthmi cell. Immunohistochemistry against GFP (in green) and against ChAT (in magenta) is shown. Arrows indicate projections to both tecta. **b.** Tectobulbar cells (with short axons) projections to the ipsilateral nucleus isthmi region. Gabaergic nucleus isthmi in magenta and glutamatergic nucleus isthmi in green. **c.** Tectobulbar cells (with long axons) projections to the ipsilateral nucleus isthmi region. **d.** Intertectal neurons projections to ipsilateral and contralateral nucleus isthmi region. **e.** Glutamatergic nucleus isthmi cells projecting to the ipsilateral thalamus (in blue, s1020t line expression). **f.** Glutamatergic nucleus isthmi cells projecting to the ipsilateral and contralateral pretectum (in blue, s1026t line expression). **g.** Glutamatergic nucleus isthmi cells projecting first to the ipsilateral tectum and then to contralateral tectum (in blue, isl2b:GFP line expression). **h.** Glutamatergic nucleus isthmi cells projecting close to the contralateral tectum and then close to ipsilateral tectum (in blue, isl2b:GFP line expression). **i.** Gabaergic nucleus isthmi cells projecting only to the ipsilateral tectum (in blue, isl2b:GFP line expression). **j.** Gabaergic nucleus isthmi cells projecting first to the ipsilateral tectum and then to the contralateral tectum (in blue, isl2b:GFP line expression). **k.** Glutamatergic NI: ipsilateral-contralateral class cells. **l.** Glutamatergic NI: contralateral-ipsilateral class cells. **m.** Glutamatergic NI: thalamus class cells. **n.** Gabaergic NI: ipsilateral-contralateral class cells. Scale bars represent 50 µm. **o.** Cellular-resolution atlas of isthmotectal circuitry. Tectal cells are shown in blue, GABAergic NI cells in magenta and glutamatergic/cholinergic NI cells in green.

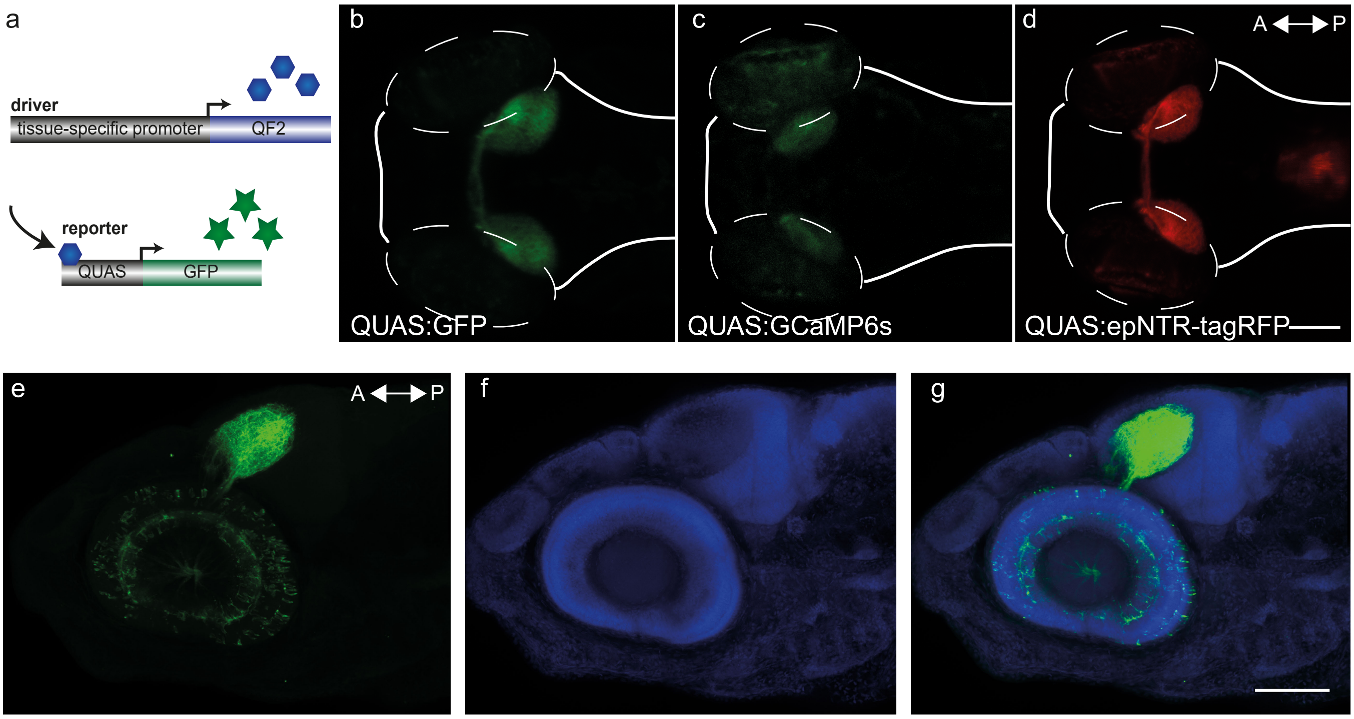
a
driver
tissue-specific promoter
QF2
reporter
QUAS
GFP
b
c
d
A
P
QUAS:GFP
QUAS:GCaMP6s
QUAS:epNTR-tagRFP
e
A
P
f
g

**Extended Figure 9.** Newly developed QF2 transgenic lines to label retinal circuits. **a.** Schematic of the QF2 system components. QF2 expressed in specific neurons activates the QUAS sequence controlling the expression of a target gene (e.g. GFP). **b.** QF2 (ath5:QF2) activation of QUAS:GFP (in green) in RGCs in a live zebrafish larvae. **c.** QF2 (ath5:QF2) activation of QUAS:GCaMP6s (in green) in RGCs in a live zebrafish larvae. **d.** QF2 (ath5:QF2) activation of QUAS:epNTR-tagRFP (in red) in RGCs in a live zebrafish larvae. **e-g**. Immunohistochemistry anti-GFP (in green) showing expression of GFP in RGC axons. DAPI staining is shown in blue. Lateral view. Scale bars represent 100 µm.

#### METHODS

##### Transgenic zebrafish lines

For the experiments in this study, we used 5-8 days post fertilisation (d.p.f.) larvae carrying mutations in the *mitfa* allele (nacre). Fish were raised on a 14h light/ 10h dark cycle at 28°C. All animal procedures conformed to the institutional guidelines set by the Max Planck Society and were approved under licenses from the regional government of Upper Bavaria (Regierung von Oberbayern).

Transgenic lines used in this study are shown in Table 1.

Table 1. List of transgenic lines

| Line | Reference |
|---|---|
| Tg(ath5:QF2)mpn405 | This study |
| Tg(QUAS:GFPcaax)mpn163 | This study |
| Tg(QUAS:GCaMP6s)mpn164 | This study |
| Tg(QUAS:epNTR-tagRFP)mpn165 | This study |
| Tg(elavl3:lyn-tagRFP)mpn404 | Dal Maschio et., 2017 |
| Tg(UAS:ChR2(H134R)-mCherry)mpn134 | Dal Maschio et al., 2017 |
| Tg(elavl3:nlsGCaMP6s)mpn400 | Dal Maschio et al., 2017 |
| Tg(gad1b:Gal4VP16) mpn155 | Förster et al., 2017 |
| Tg(lhx9:Gal4VP16) mpn203 | Förster et al., 2017 |
| Tg(chata:Gal4VP16) mpn202 | Förster et al., 2017 |
| SAGFF(LF)81C | Sato et al., 2015 |
| Tg(UAS:nfsb-mCherry)c264 | Davison et al., 2007 |
| Tg(-7atoh7:GAL4-VP16) s1992tTg | Del Bene et., 2010 |
| Tg(gad1b: loxP-DsRed-loxP-GFP) | Satou et al., 2013 |
| Tg(vglut2a:loxP-DsRedloxP-GFP) | Satou et al., 2013 |
| Tg(UAS:GCaMP6s)mpn101 | Thiele et al., 2014 |
| *Tg(UAS:Dendra-kras)s1998t* | Arrenberg et al., 2009 |
| Tg(elavl3:GCaMP6s) a13203 | Kim et al., 2017 |
| Tg(isl2b:Gal4-VP16)zc65 | Fujimoto et al., 2011 |

##### Q genetic system

A pTol2-(5x)QUAS-e1b:EcoRV-polyA;cmlc2:mCherry vector was generated using oligo synthesised QUAS promoter sequences[1]. A GCaMP6s fragment[2] was inserted into the EcoRV-linearised vector using traditional restriction-ligation cloning. A similar approach was used to generate an eNTR-TagRFP[3] and GFPcaax[4] version. To generate the Tg(QUAS) lines, transposase mRNA (50ng/µl) and the construct (25ng/µl) were co-injected into TL/nacre zebrafish eggs. Injected fish were sorted based on heart expression of the transgenesis marker cmlc2:mCherry and raised into adulthood. Founders were obtained by outcrosses to TL/nacre.

The QF2 coding sequence was obtained from Addgene Plasmid #61312 [5]. To establish the driver line Tg(ath5:QF2), a QF2-polyA; FRT-KAN-FRT;cmlc2:Cerulean fragment was PCR amplified and inserted into the ath5 BAC clone DKEY-111E19 using BAC recombineering as described previously[4]. The resulting Tol2-ath5:QF2 BAC (100ng/µl) was co-injected with transposase mRNA (50ng/µl) into TL/nacre zebrafish eggs. cmlc2:Cerulean expressing larvae were raised and identified as founders by an outcross to Tg(QUAS:GCaMP6s) or Tg(QUAS: eNTR-TagRFP) fish.

**Visual competition behavioural assay**

We designed a setup to project looming stimuli to 9 individual zebrafish larvae simultaneously via a screen below the animals. To this end, we adapted a previously described virtual reality setup which allows real-time tracking and presentation of arbitrary visual stimuli at animal-centric positions[6]. Animals were monitored individually in shallow glass dishes of 10 cm diameter separated by opaque walls to prevent visual contact. The dishes rested on a projection film for visual stimulation, confining the animals' distance to the screen to approximately between 3 mm and 13 mm by the rounded bottom of the dish and the surface of the water. To minimise stimulus distortion due to refraction at the air-glass-water interfaces, we submerged the projection film and glass dishes in water. Animals were recorded at 30 fps with cameras using the CMV4000 sCMOS chip (IDS UI-3370CP-NIR or PointGrey Grasshopper GS3-U3-41C6NIR-C) at a resolution of 2048x2048 pixels. We used a 25 mm lens (Edmund Optics Nr. 86-572) at a distance of 800 mm resulting in an image resolution of 150 um/pixel. Visual stimuli were projected onto the projection film from underneath via a cold mirror. Diffuse Infrared illumination for imaging was provided from below. Visible light and stimuli were provided to the fish via the projector but blocked before the camera by an IR band-pass filter. Image acquisition at 30 fps, real-time processing and stimulus generation were performed on a Desktop PC running Bonsai[7]. Briefly, each camera frame was background subtracted and a threshold was applied to isolate animals against the background. Next, contours were extracted to compute the centre of mass and orientation of each animal.

Based on animal positions and a stimulus property file, we generated animal-centric visual stimuli using custom Python scripts in Bonsai to control OpenGL drawing routines. Stimuli were dots (for black, RGB value (0,0,0)) on white background (RGB value (255,255,255)) unless noted otherwise. Dot size was a multiple of projected pixel size (1 px was 0.47 mm side length). Loom stimuli were presented as stationary dots expanding for 500 milliseconds (15 frames) with a linear increase in diameter.
Stimuli were presented 1cm from the fish at angles of 45°, 90°, 135°, 180°, 225°, 270° or 315° relative to relative to the animals' centre of mass and orientation at the beginning of the stimulus. Loom stimuli were presented once per minute. A moving grating was presented for 20 seconds ending 20 seconds before the presentation of the next loom stimulus to drive larvae towards the centre of the dishes. At each frame, animal and stimulus parameters were streamed to a text file for offline analysis. The program also stored the video data after background subtraction into an xVid compressed .avi file via FFmpeg (ffmpeg.org) for later inspection. Camera and projector were aligned using a separate Bonsai routine once per day as described previously [6].
Animals were tested at 5-8 dpf in fish water at room temperature (22-25°C). Before behaviour testing, animals were kept in a petri dish floating above a fully lit portion of the projection screen to allow habituation to light and temperature conditions of the experiment. Animals were analyzed for 60 to 180 minutes. The order of different stimuli was randomised for each group of 9 animals.

**Prey capture experiments**

Prey capture experiments were performed in a custom-built square chamber (15 x 15 mm, 5 mm deep) with walls made from 2% agarose and a glass coverslip placed over the top. Individual larvae were introduced to the behaviour chamber with a drop of dense paramecia culture (*Paramecium multimicronucleatum*, Carolina Biological Supply Company, Burlington, NC). The setup was lit from below with an IR LEDs light source, and larvae were filmed for 20 minutes at 500fps with a high-speed camera (Photonfocus MV1-D1312-160-CL, Switzerland). The analysis was performed offline with custom-written Python code. We extracted the outline of the fish from each frame by finding the largest contour following background subtraction and thresholding. A second threshold was then set to extract contours of the eyes and swim bladder. We used the image moments of these contours to calculate the angle of each eye. Eye convergence in each frame was calculated as the difference between the eye angles, with positive values corresponding to eye convergence, zero corresponding to eyes parallel, and negative angles corresponding to eye divergence. For each fish, we defined the threshold for prey engagement as the anti-mode of the bimodal distribution of eye convergence angles across all frames and defined the prey capture score as the proportion of time the fish spent engaged in prey capture.

**Data analysis of behaviour experiments**

Exported text files containing behavioural data and stimulus parameters were analyzed offline using custom-written Python scripts. We classified responses as escapes if the distance to the original position at the end of the expansion time of the stimuli (500ms, after 15 frames) was at least 5 mm (approximately one fish body length). Distance from the initial position was defined as the Euclidean distance from the origin to a point in the x-y plane after 500ms (end of stimuli expansion). The distance modulation of escape behaviours is in agreement with a previous study[8] and was used as an indication of stimulus strength. Circular behavioural data statistics was performed with the python version of pycircstat[9] (https://github.com/circstat/pycircstat).

**Modelling**

All models were implemented in Python, using numpy and scipy libraries. All models are based on repeated random sampling, where one stimulus response from an S1 trial and one stimulus response from an S2 trial are combined. The repetition of this sampling procedure generates a distribution of combined responses. The averaging model combines the pair of responses by taking the vector average of the response angle. In agreement with the reduced amount of backward responses, we implemented a mechanism to reduce the prevalence of such escapes in our model by redistributing backwards swims to other headings. The winner-take-all model chooses randomly between the S1 response and the S2 response (effectively adding the S1 and S2 response distribution). The mixture model implements a random assortment between the winner-take-all model (with probability p) and the average model (with probability p-1). Distributions are plotted using a kernel density estimate (KDE) plot, with a von Mises (circularised) distribution. To compare the similarity of distributions, a circularised version of the energy distance metric was used.

#### Imaging

Calcium imaging: Zebrafish larvae were embedded in 2.5% low melting point agarose (Invitrogen). Visual stimuli were generated using custom Psychopy2 scripts[10] and consisted of black looming stimuli. For prey competition (Extended Fig.4a) movies from recordings of real paramecia have been binarised and scaled, keeping important parameters such as kinetics and size in agreement with [11]. Visual stimuli were projected onto a white diffusive screen using the red channel of a LED projector (LG, Model No. PA72G) from the side (animal distance to the screen was approximately 4cm) and a DLP® LightCrafter™ 4500 development module from the bottom (animal distance to the screen was approximately 1cm). Size of the stimuli (in degrees of visual angle) was adjusted taking into account the size of the projection and distance to the fish using Psychopy2 Monitor Center. For monocular stimulation, we presented both from the side and bottom. Results were similar for both conditions. The minimum distance between stimuli resulting in non-overlapping receptive fields and suppression was determined in pilot experiments to be at least 30 degrees in visual space.

The 2P microscope used for imaging and holographic optogenetics is based on a modified Femtonics 3DRC (Femtonics, Hungary) driven by a Ti:Sapphire source (Chameleon Ultra II, Coherent) (see [12]). An electrically tunable lens (ETL, Optotune, EL-10-30-Ci-IR-LD-MV) was used to enable fast remote refocusing.

For Extended Fig.3a we used a remote-Z-scanning module with a resonant 2P microscope. The module consists of a second objective and a piezo-modulated mirror, which allows us to shift between conjugated focal planes in the fish brain with high frequency.

#### Computer Generated Holography (CGH) photostimulation

The approach used is similar to Dal Maschio et al 2017 [12]. Optogenetic stimulation of ChR2 positive neurons was performed with 920 nm excitation with a total duration of 1000ms (photostimulation started 500ms before the visual stimulation and ended 500ms later at the end of visual stimulation). Visual stimulation consisted of a single looming stimulus (total duration of expansion of 500ms, 60°/s expansion rate). Imaging was performed simultaneously with GCaMP6s at 1,020 nm.

**Genetic ablation of neurons**

Larvae expressing Tg(UAS:nfsb-mCherry)c264, were treated with 7,5mM metronidazole (MTZ, Sigma Aldrich) in fish water containing 0.2% DMSO, typically for at a minimum of 8h in a light protected chamber. MTZ solution was washed and larvae were allowed to recover for at least 12h before imaging or behavioural experiments were performed.

**Imaging analysis**

Imaging analysis was performed with custom-written Python scripts. A regressor-based pixel-wise analysis of the imaging data was performed similarly to [13]. For Fig. 2i a linear regression was used (Python scipy.stats.linregress). For ROI analysis, a linear regression approach was used (Python scikit-learn) similar to [14]. We used the ordinary least-squares linear regression, y = a + b0x0 + b1x1 + e... (y representing the functional response, a representing the Y-intercept, b the coefficients (slope), x the regressors and e the residual error) to select ROIs. The coefficients of determination (R2), were calculated using the sklearn.linear_model.LinearRegression method. R2 was used to set a threshold removing ROIs with activity not locked to stimulus presentation (spontaneously active). The estimated coefficients for the linear regression problem were used to set a second threshold that selects ROIs fitting to the regressors used (time series set to zero for all time points except the time points of visual stimulation). For quantification of holographic optogenetic activation effects, we generated a control distribution by shuffling the labels of trials with visual alone and trials with visual combined with optogenetic stimulation. This approach led to a normal distribution with a strong peak at around zero, used to set thresholds considered for quantification of enhanced and suppressed ROIs.

**Quantification and Statistical Analysis**

For statistical tests, we used the Python SciPy library and GraphPad Prism version 7 for Windows. All statistical tests used were two-tailed tests. For imaging experiments, we preferentially selected fish with strong expression. All error bars used are mentioned in figure legends.

**Data and Software Availability**

The datasets and custom software that support the findings of this study will be made available upon request. The code used to model the behavioural data will be made available at Bitbucket (link active upon publication).

**Single-cell reconstructions**

For some of the single neuron labelling, the transgenic fish line *Tg(brn3c:GAL4, UAS:gap43-GFP)s318t* (BGUG) was crossed to *Tg(lhx9:Gal4)mpn203* line similar to [14]. This approach could not be used with other lines (e.g. *Tg(gad1b:Gal4VP16) mpn155)* possibly due to low expression level. Also with the *Tg(lhx9:Gal4)mpn203* line we could rarely label single cells for some subtypes possibly due to clonal effects hindering sparse labelling. To overcome this problem, we devised another method to achieve sparse labelling by co-injecting a plasmid with a heat shock promoter expressing Cre (hsp70l:cre) together with a UAS:Brainbow plasmid (UAS:Brb1.0L [15]). These constructs were co-injected with Tol2 mRNA into *Tg(lhx9:Gal4)mpn203 or Tg(lhx9:Gal4)mpn203* ; Tg(UAS:nfsb-mCherry)c264 embryos. By calibrating the heat shock duration (heat shock in a water bath at 37°C for 5-45 min), the EYFP fluorescence from the UAS:Brainbow construct could be used to label single cells. Constructs were pressure injected at a concentration of 25–50 ng/µL into 1–4 cell-stage embryos. Larvae were screened using a confocal microscope for single labelled projection neurons and positive larvae were used to record a high-resolution (1024 x 1024 pixels) confocal stack

**Confocal imaging and anatomical reconstruction of neurons**Before acquiring confocal stacks, fish were anaesthetised with 0.02% tricaine. For single-cell reconstructions and generation of the brain, atlas imaging was performed as described previously[14]. The collected neurons were then traced using the software neuTube (Build1.0z) and confirmed by at least two independent tracers. For live-imaging rainbow experiments, no reference channel was available. However, in the YFP-channel the signal was strong not only for the single neurons but also for the auto-fluorescence of the skin. We took advantage of this and registered the whole-brain YFP-stacks to a standard brain of the skin auto-fluorescence using the Computational Morphometry toolkit (CMTK - https://www.nitrc.org/projects/cmtk/, [16]) with the following settings: -awr 01 -T30 -X52 -C8 -G120 -R3 -A'–accuracy 0.8' -W'–accuracy 0.8'. This standard brain was generated by registering the red and green channel of 150 fish expressing *elavl3:lyn-tagRFP* to the standard brain as described in [14]. The registered green channel of these fish was then averaged to obtain a standard brain of the skin auto-fluorescence. In experiments using fixed animals, fish were stained against GFP for single neurons and synapsin as a whole brain marker and registered to the fixed synapsin standard brain as described in [17]. The skin-registered neurons were then bridged to the synapsin standard brain using the *elavl3:lyn-tagRFP* channel. All reconstructed neurons were visualised in their common coordinate system (synapsin) using custom-written Python code employing the Mayavi library (Figure 4 a-g) or the single-neurite tracer ImageJ plugin (Extended Fig. 8). Standard brains (s1026t, s1020t, isl2b:GFP$^{zc7Tg}$) used in Extended Fig.8 are part of the brain atlas in [17].

**In Situ Hybridisation and Immunohistochemistry**

Stainings were performed according to [18]. Antibodies used were anti-TH (MAB318, Millipore), anti-REELIN (40-189, Millipore), anti-GFP(632380, Takara Bio Clontech), anti-GFP(A10262, Molecular Probes) and anti-CHAT (AB144P, Millipore). Riboprobes for *adcyap1a*, *nnos1* and *lhx9*, were generated from cDNA and subcloned into the TOPO vector (pCR2.1-TOPO, Invitrogen). Sense probes were used as a negative control for newly cloned probes. Riboprobes for *gad67* and *trh*[19] were a kind gift of Wolfgang Driever. For Extended Fig. 9, DAPI (28718-90-3, Sigma) was used.

**References Extended Data**

17. A Cellular-Resolution Atlas of the Larval Zebrafish Brain by Michael Kunst, Eva Laurell, Nouwar Mokayes, Anna Kramer, Fumi Kubo, Antonio M. Fernandes, Dominique Förster, Marco Dal Maschio, Herwig Baier :: SSRN. Available at: https://papers.ssrn.com/sol3/papers.cfm?abstract_id=3257346.

18. Fernandes, A. M. et al. Deep brain photoreceptors control light-seeking behavior in zebrafish larvae. Curr. Biol. 22, 2042–2047 (2012).

19. Löhr, H., Ryu, S. & Driever, W. Zebrafish diencephalic A11-related dopaminergic neurons share a conserved transcriptional network with neuroendocrine cell lineages. Development 136, 1007–1017 (2009).
